# A field-wide assessment of differential expression profiling by high throughput sequencing reveals widespread bias

**DOI:** 10.1101/2021.01.04.424681

**Authors:** Taavi Päll, Hannes Luidalepp, Tanel Tenson, Ülo Maiväli

## Abstract

Here we assess inferential quality in the field of differential expression profiling by high throughput sequencing, based on analysis of datasets submitted 2008-2020 to the NCBI GEO data repository. We take advantage of the parallel differential expression testing over thousands of genes, whereby each experiment leads to a large set of p values, the distribution of which can indicate the validity of assumptions behind the test. Moreover, from a well-behaved p value set *π*_0_, the fraction of genes that are not differentially expressed, can be estimated. We found that only 25% of experiments resulted in theoretically expected p value histogram shapes, although there is a marked improvement over time. Uniform p value histogram shapes, indicative of < 100 true effects, were extremely few. Furthermore, although many HT-seq workflows assume that most genes are not differentially expressed we found 37% of experiments to have *π*_0_-s of less than 0.5, as if most genes changed their expression level. Restricting our analysis to studies involving cancer or transcription factors, expected to lead to real changes in expression of many genes, did not result in meaningfully different distributions of *π*_0_-s. Both the fractions of different p value histogram types and the *π*_0_ values are strongly associated with the differential expression analysis program used by the original authors. While we could double the proportion of theoretically expected p value distributions by removing low-count features from analysis, this treatment did not remove the association with the analysis program. Taken together, our results indicate widespread bias in differential expression profiling field.

## Introduction

Over the past decade a feeling that there is a crisis in experimental science has increasingly permeated the thinking of methodologists, captains of industry, working scientists, and even the lay public [1–6]. This manifests in poor statistical power to find true effects [7,8], in poor reproducibility (defined as getting identical results when reanalysing the original data by the original analytic workflow), and in poor replicability (defined as getting similar results after repeating the entire experiment) of the results [9]. The proposed reasons behind the crisis include sloppy experimentation, selective publishing, perverse incentives, difficult-to-run experimental systems, insufficient sample sizes, over-reliance on null hypothesis testing, and much-too-flexible analytic designs combined with hypothesis-free study of massively parallel measurements [10–15]. Although there have been attempts at assessing experimental quality through replication of experiments, mostly in psychology, prohibitive costs, and theoretical shortcomings in analysing concordance in experimental results have encumbered this approach in biomedicine [16–18]. Another way to assess the large-scale quality of a science is to employ surrogate measures for quality that can be more easily obtained than full replication of a study. The most often used such measure is technical reproducibility, which involves checking for availability of code and/or running the original analysis code on the original data. Although the evidence-base for reproducibility is still sketchy, it seems to be well below 50% in several fields of biomedicine [18]. However, as there are many reasons, why a successful reproduction might not indicate a good quality of the original study, or why an unsuccessful reproduction may not indicate a bad quality of the original study, the criterion of reproducibility is clearly insufficient. Another proxy for quality can be found in published p values, especially the distribution of p values [19]. In a pioneering work Jager and Leek extracted ca. 5000 statistically significant p values from abstracts of leading medical journals and pooled them to formally estimate, from the shape of the ensuing p value distribution, the Science-Wide False Discovery Rate or SWFDR as 14% [20]. However, as this estimate rather implausibly presupposes that the original p values were calculated uniformly correctly, and that unbiased sets of significant p values were obtained from the abstracts, they subsequently revised their estimate of SWFDR upwards, as “likely not >50%” [21]. For observational medical studies, by a different method, a plausible estimate for field-wide FDR was found to be somewhere between 55% and 85%, depending on the study type [22]. While our work uses published p values as evidence for field-wide quality and presupposes access to unbiased full sets of unadjusted p values, it does not pool the p values across studies, nor does it assume that they were correctly calculated. In fact, we assume the opposite and do a study-by-study analysis of the quality of calculation of p values. This makes the quality of the p value a proxy for the quality of the experiment and/or of the scientific inferences based on these p values. We do not see our estimate of the fraction of poorly calculated p values as a formal quality metric, but merely hope that by this measure we can shed some light into the overall quality of a field. However, we chose the field, whose quality to assess, so as to maximize the weight p values could have on scientific inferences. We concentrate on field of differential expression profiling studies using high throughput sequencing (DE HT-seq, mostly RNA-seq) for two reasons. HT-seq has become the gold standard for whole transcriptome gene expression quantification, both in research and in clinical applications [23]. And secondly, due to the massively parallel testing in individual studies of tens of thousands of features per experiment, we can access study-wide lists of p values. From the shapes of histograms of p values, we can identify the experiments where p values were calculated apparently correctly, and from these studies we can estimate the study-wise relative frequencies of true null hypotheses (the *π*_0_-s). Also, we believe that the very nature of the DE HT-seq field, where a single biological experiment entails comparing the expression levels of about 20,000 different features (e.g. RNA-s) on average, predicates that the quality of data analysis, and specifically statistical inference based on p values (directly, or indirectly through FDR) must play a decisive part in scientific inference. Simply, one cannot analyse a DE HT-seq experiment intuitively, without resorting to formal statistical inference. Therefore, quality problems of statistical analysis would very likely directly and substantially impact the quality of science. Thus we use the quality of statistical analysis as a proxy for the quality of science, with the understanding that this proxy may work better for modern data-intensive fields, where scientist’s intuition has a comparatively smaller role to play.

## Results

### Data mining

We queried the NCBI GEO database for “expression profiling by high throughput sequencing” (for the exact query string, see Methods), retrieving 43,610 datasets (GEO series) from 2006, when first HT-seq dataset was submitted to GEO, to Dec-31, 2020. The number of yearly new HT-seq submissions increased from 1 in 2006 to 11,604 by 2020, making up 26.6% of all GEO submissions in 2020. First, we filtered the GEO series containing supplementary processed data files. NCBI GEO database submissions follow MINSEQE guidelines [24]. Processed data are a required part of GEO submissions, defined as the data on which the conclusions in the related manuscript are based. The format of processed data files submitted to GEO is not standardized, but in case of expression profiling such files include, but are not limited to, quantitative data for features of interest, e.g. mRNA, in tabular format. Sequence read alignment files and coordinates (SAM, BAM, and BED) are not considered as processed data by GEO. According to our analysis the 43,610 GEO series contained 84,036 supplementary data files, including RAW.tar archives. After unpacking RAW.tar files, we programmatically attempted to import 647,092 files, resulting in 336,602 (52%) successfully imported files, whereas 252,685 (39%) files were not imported because they were either SAM, BAM, BED, or in other formats most probably not containing p values. We failed to import 57,805 (8.9%) files for various reasons, mostly because of text encoding issues and failure to identify column delimiters. According to GEO submission requirements, the processed data files may contain raw counts of sequencing reads, and/or normalized abundance measurements. Therefore, a valid processed data submission may or may not contain lists of p values. We identified p values from 4,616 GEO series, from which we extracted 14,813 unique unadjusted p value sets. While the mean number of p value sets, each set corresponding to a separate experiment, per 4,616 GEO submissions was 3.21 (max 276), 46% of submissions contained a single p value set and 76% contained 1-3 p value sets. For further analysis we randomly selected one p value set per GEO series.

### P value histograms

We algorithmically classified the p value histograms into five classes (See Methods for details and Fig. 1A for representative examples) [25]. The “Uniform” class contains flat p value histograms indicating no true effects (at the sample sizes used to calculate these p values). The “Anti-Conservative” class contains otherwise flat histograms that contain a spike near zero. The “Conservative” class contains histograms that have a distinct spike close to one. The “Bimodal” histograms have two peaks, one at either end. The class “Other” contains a panoply of malformed histogram shapes (humps in the middle, gradual increases towards one, spiky histograms, etc.). The “Uniform” and “Anti-Conservative” histograms are the theoretically expected shapes of p value histograms.

**Fig 1.**
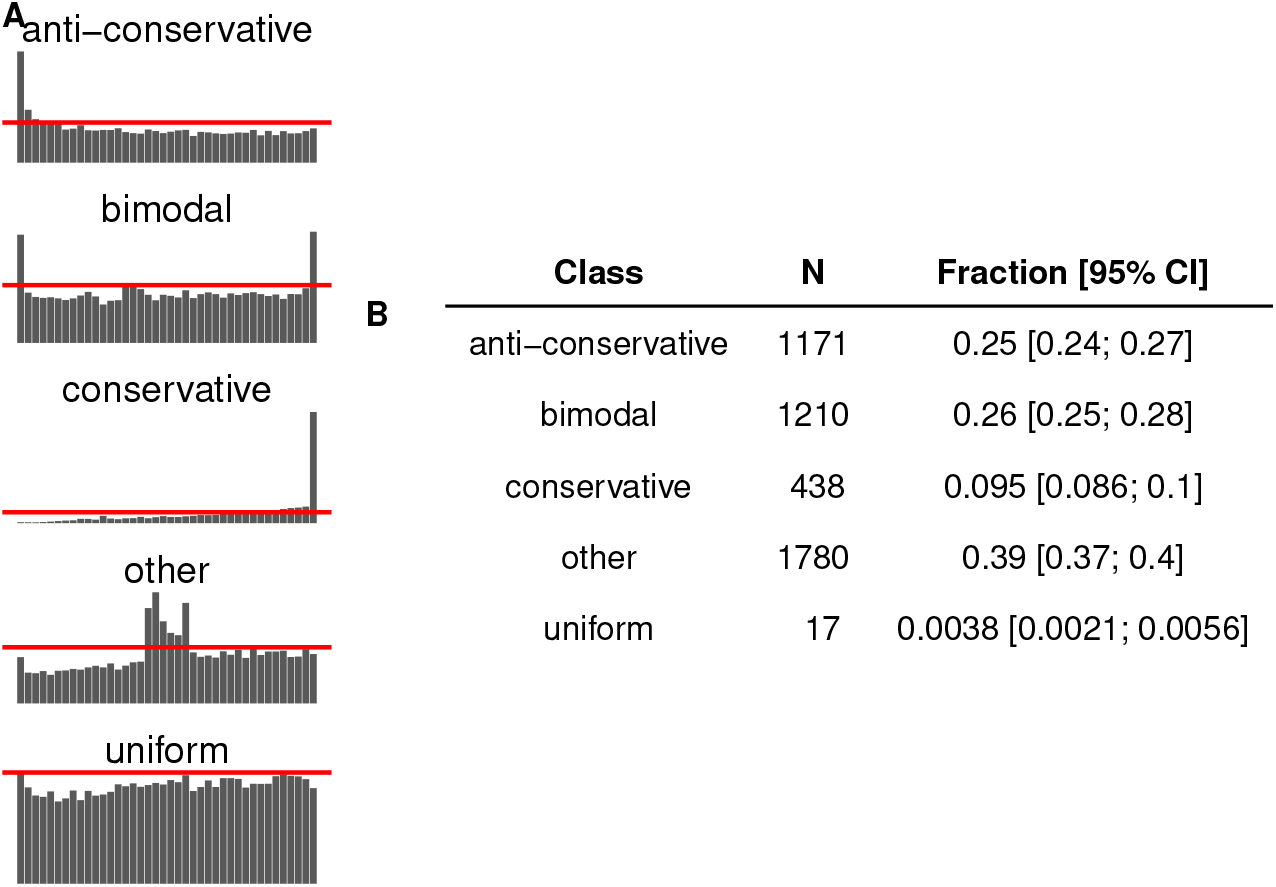
Classes of p value histograms. (**A**) Examples of p value histogram classes. Red lines denote the algorithmic threshold used for separating p value histograms into discrete classes. (**B**) Summary of p value histograms identified from GEO supplementary files. One p value set was randomly sampled from each GEO series where p values were identified. N = 4,616. 95% CI are credible intervals. The model object related to panel B can be downloaded from https://gin.g-node.org/tpall/geo-htseq-paper/raw/26619a4b74aa3781ac6a244edcc24e0ad6eb064b/models/Class_1.rds.

We found that overall, 25% of the histograms fall into anti-conservative class, 9.5% were conservative, 26% bimodal and 39% fell into class “other” (Fig. 1B). Only 17 of the 4,616 histograms were classified as “uniform.” Median number of features in our sample was 20,954. Interestingly, there is an apparent leftward shift in the peak of distribution of features in anti-conservative histograms, as compared to histograms with all other shapes, suggesting different data pre-processing for datasets resulting in anti-conservative histograms (S1 Fig.). Logistic regression reveals a clear trend for increasing proportion of anti-conservative histograms, starting from <10% in 2010 and topping 30% in 2020 (S2 Fig.). Hierarchical modelling indicates that all differential expression (DE) analysis tools and sequencing platforms exhibit similar temporal increases of anti-conservative p value histograms (S3-4 Fig.). Multinomial hierarchical logistic regression further demonstrated that the increase in the fraction of anti-conservative histograms is accomplished by decreases mostly in the class “other”, irrespective of the DE analysis tool (Fig. 2A).

**Fig 2.**
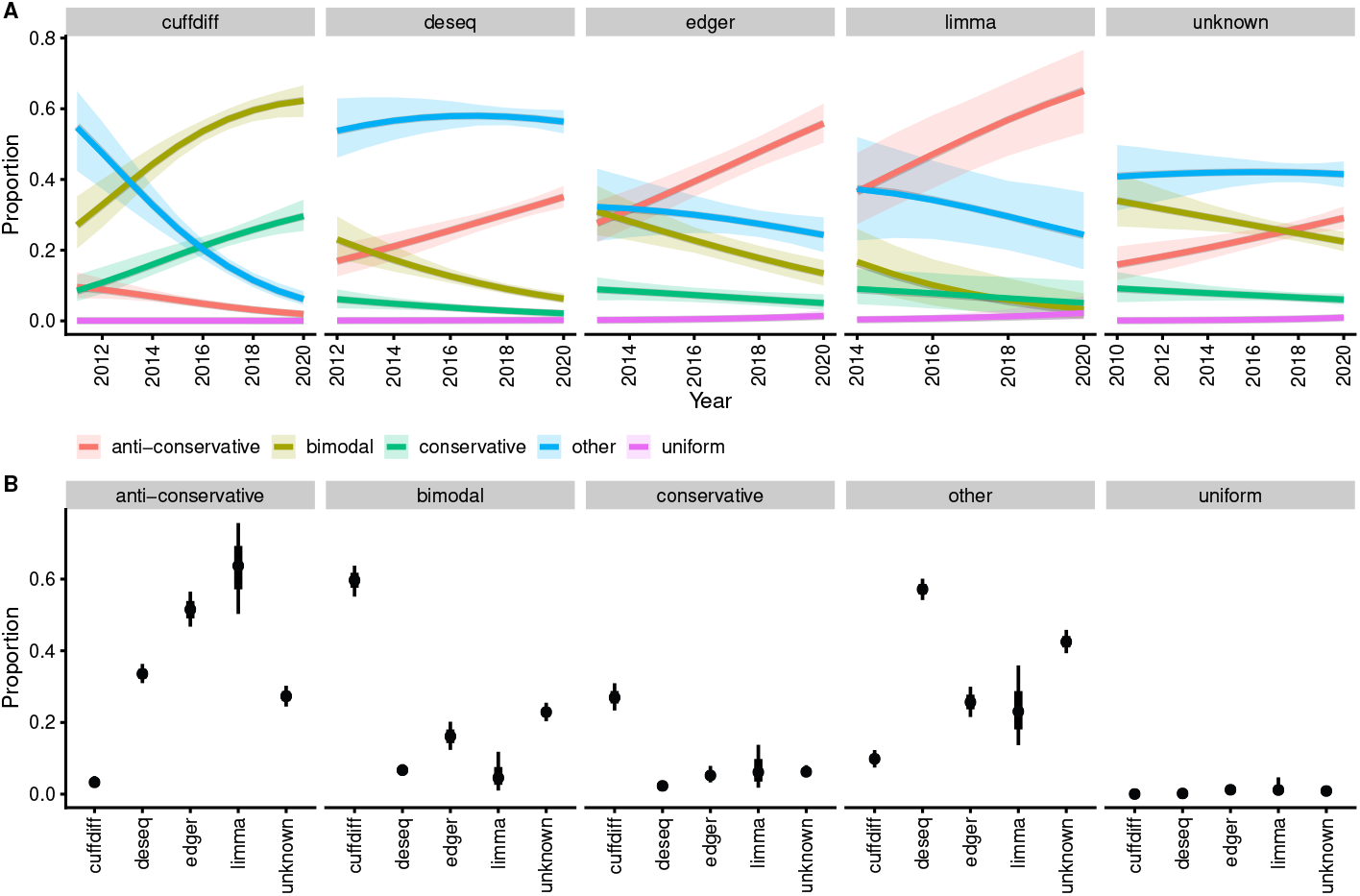
Association of the p value histogram class with differential expression analysis tool. (**A**) The increase in the proportion of anti-conservative histograms is accompanied by decreases mostly in the class “other,” irrespective of the DE analysis tool. Lines denote best fit of the model [class ∼ year + (year | de_tool), categorical likelihood]. Shaded areas denote 95% credible regions. N = 4,616. (**B**) Association of p value histogram type with DE analysis tool; data is restricted to 2018-2020 GEO submissions. Points denote best fit of the model [class ∼ de_tool, categorical likelihood]. Thick and thin lines denote 66% and 95% credible intervals, respectively. N = 2,930. The model object related to panel A can be downloaded from https://gin.g-node.org/tpall/geo-htseq-paper/raw/26619a4b74aa3781ac6a244edcc24e0ad6eb064b/models/Class_year_year_detool_year.rds. The model object related to panel B can be downloaded from https://gin.g-node.org/tpall/geo-htseq-paper/raw/26619a4b74aa3781ac6a244edcc24e0ad6eb064b/models/Class_detool_2018up.rds.

This positive temporal trend in anti-conservative p value histograms suggests improving quality of the DE HT-seq field. Rather surprisingly, Fig. 2A also indicates that different DE analysis tools are associated with very different proportions of p value histogram classes, suggesting that quality of p value calculation, and therefore, quality of scientific inferences based on these p values depends on DE analysis tool. We further tested this conjecture in a simplified model, restricting our analysis to 2018-2020, the final years in our dataset (Fig. 2B). As no single DE analysis tool dominates the field – cuffdiff 23%, deseq 33%, edgeR 13%, limma 2%, unknown 29% (see S5 Fig. for temporal trends) –, a state of affairs where proportions of different p value histogram classes do not substantially differ between analysis tools would indicate lack of DE analysis tool generated bias to the results. However, we found by multinomial regression that all p value histogram classes, except “uniform,” which is largely unpopulated, depend strongly on the DE analysis tool used to calculate the p values (Fig. 2B). This is confirmed by modelling the frequency of the anti-conservative p value histograms in binomial logistic regression (panel A in S6 Fig.). Using the whole dataset of 14,813 p value histograms – as a check for robustness of results – or adjusting the analysis for GEO publication year, of taxon (human, mouse, and pooled other), of the RNA source or sequencing platform – as a check for possible confounding – does not change this conclusion (panel B-E in S6 Fig.). The lack of confounding in our results allow a causal interpretation, indicating that DE analysis tools bias the analysis of HT-seq experiments [26].

### Proportion of true null hypotheses

To further enquire into DE analysis tool-driven bias we estimated from user-submitted p values the fraction of true null effects (the *π*_0_) for each HT-seq experiment. As non-anti-conservative sets of p values (excepting the “uniform”) indicate problems during the respective experiments and/or data analyses, we only calculated the *π*_0_ for datasets with anti-conservative and uniform p value distributions (N = 1,188). Nevertheless, the *π*_0_-s show an extremely wide distribution, ranging from 0.999 to 0.02. Remarkably, 37% of the *π*_0_ values are smaller than 0.5, meaning that in those experiments over half of the features (e.g. mRNA-s) are estimated to change their expression levels upon experimental treatment (Fig. 3A). Conversely, only 23% of *π*_0_-s exceed 0.8, and 9.9% exceed 0.9. Furthermore, the peak of the *π*_0_ distribution is not near 1, as might be expected from experimental design considerations, but there is a wide peak between 0.5 and 0.8 (median and mean *π*_0_-s are both at 0.59). Depending on the DE analysis tool, the median *π*_0_-s range over 20 percentage points, from 0.5 to 0.7 (Fig. 3B). Using the whole dataset confirms the robustness of this analysis, also producing narrower credible intervals, due to larger sample (N = 3,898) (S7 Fig. A). In terms of experimental design, to get a low *π*_0_ an experiment would have to change the expression of most genes under study. A major source of such experiments would be comparisons of different cancer cell lines/tissues, where *π*_0_=0.4 could be considered a reasonable outcome [27]. We therefore compared the *π*_0_-s coming from GEO DE HT-seq submissions related to search terms “neoplasms” or “cancer” (for exact query string, please see Methods) with all other, non-cancer, submissions. There is very little difference in the means and standard deviations of *π*_0_-s for cancer and non-cancer experiments (0.58 (0.22) and 0.59 (0.24), respectively) (Fig. 4C). Filtering by studies mentioning transcription factor led to very similar results (Fig. 4D), suggesting that the wide dispersion of *π*_0_-s is not caused by intentional experimental designs. Also, studies involving cancer and TFs resulted in anti-conservative or uniform p value distributions with similar probabilities to non-cancer/non-TF studies (risk ratios with 95% CI are 0.95 (0.83; 1.07) and 0.99 (0.74; 1.28), respectively). In addition, controlling for time, taxon, or sequencing platform, did not substantially change the association of DE analysis tools with the *π*_0_-s (S7 Fig. B-E). Recalculating the *π*_0_-s with a different algorithm [28] did not change these conclusions (data not shown). As there is a strong association between both *π*_0_ (S8-11 Fig.) and proportion of anti-conservative p value histograms (S12-15 Fig.) with DE analysis tool (Fig 2, Fig 3), we further checked for, and failed to see, similar associations with variables from raw sequence metadata, such as the sequencing platform, library preparation strategies, library sequencing strategies, library selection strategies, and library layout (single or paired). These negative results support the conjecture of specificity of the associations with DE analysis tools.

**Fig 3.**
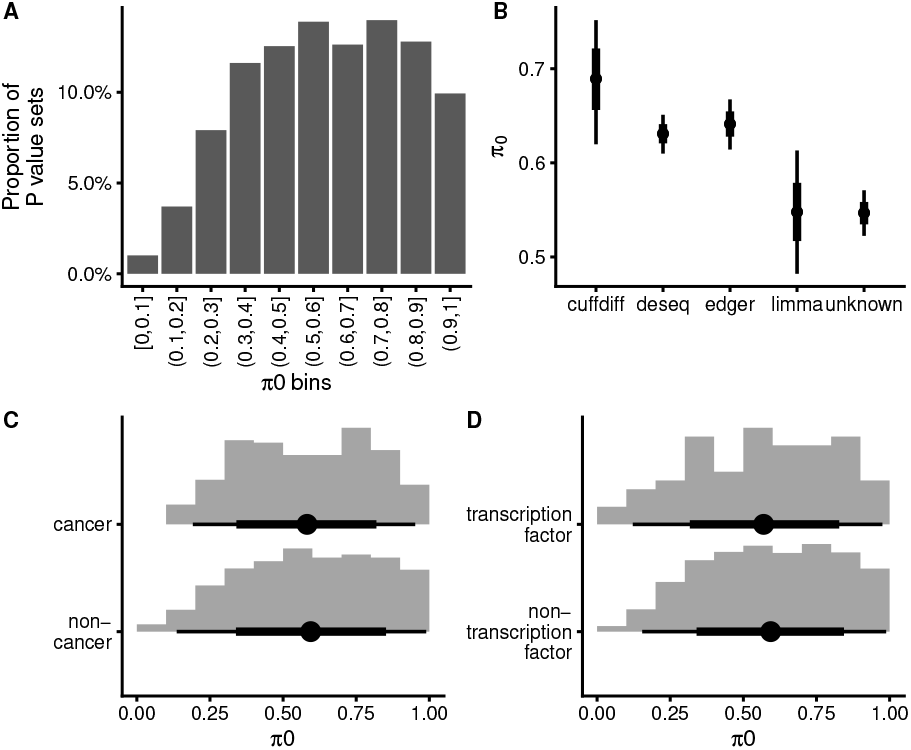
Association of the proportion of true null effects (*π*0) with DE analysis tool. (**A**) Histogram of *π*0 values estimated from anti-conservative and uniform p value sets. N = 1,188. (**B**) Robust linear model [pi0 ∼ de_tool, beta likelihood] indicates association of *π*0 with DE analysis tool. Points denote best estimates for the mean *π*0 and thick and thin lines denote 66% and 95% credible intervals, respectively. N = 1,188. (**C**) Histogram of *π*0 values in GEO cancer studies compared to non-cancer studies. (**D**) Histogram of *π*0 values in GEO transcription factor studies compared to non-TF studies. Point denotes mean, thick- and thin lines denote 66% and 95% quantile interval, respectively. The model object related to panel B can be downloaded from https://gin.g-node.org/tpall/geo-htseq-paper/raw/26619a4b74aa3781ac6a244edcc24e0ad6eb064b/models/pi0_detool_sample.rds.

**Fig 4.**
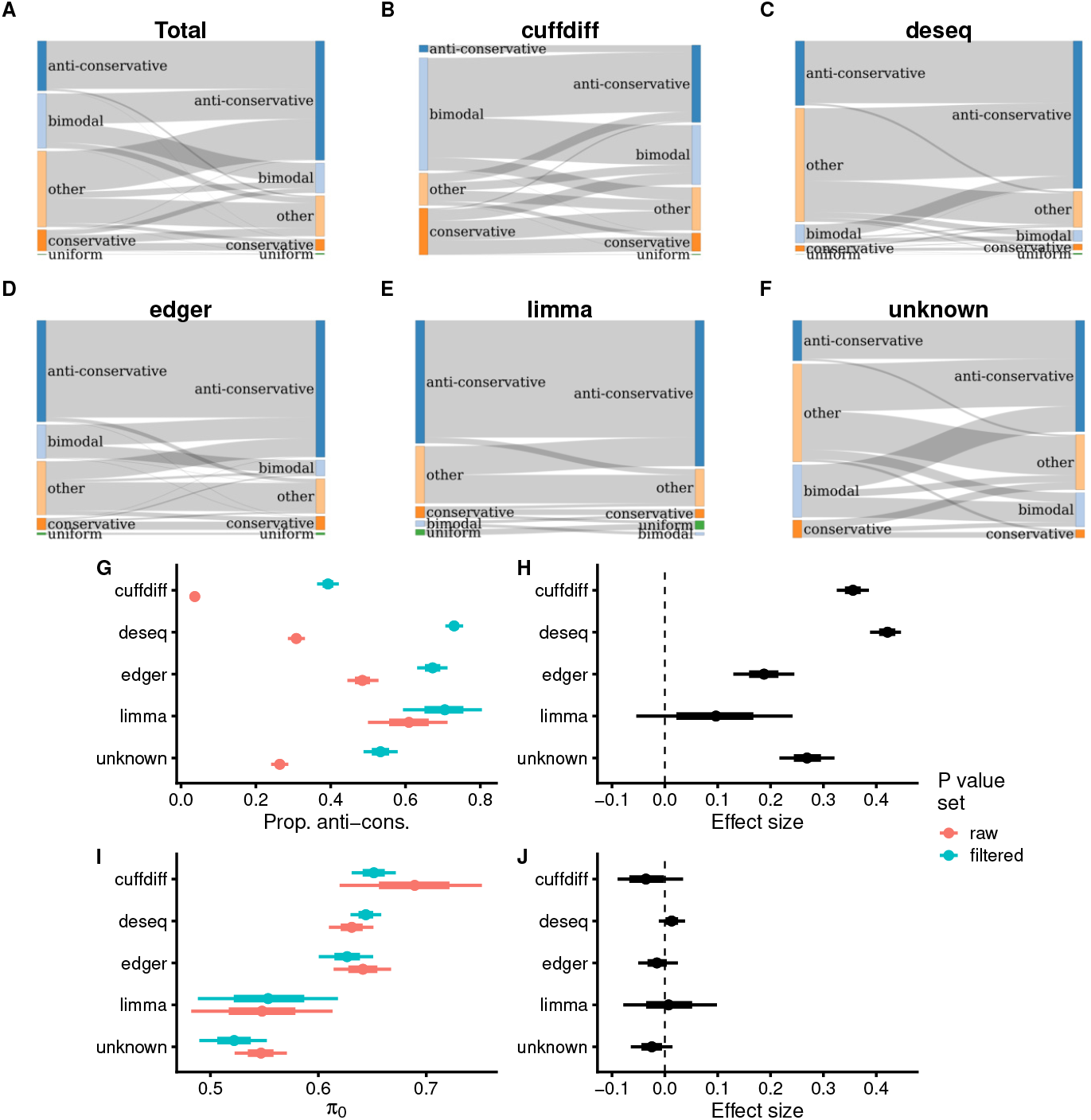
Removal of low-count features results in increasing proportion of anti-conservative p value histograms. **A-F**. Flow chart of transformation of p value histogram shape. Ribbon size is linearly proportional to the number of p value sets that change their distributional class. Only the 3,550 experiments that could be subjected to this treatment are depicted. (**A**) Full data, N = 3,550. (**B**) The subset where the p values were calculated with cuffdiff, N= 1,137. (**C**) The subset where the p values were calculated with DESeq/DESeq2, N= 1,393. (**D**) The subset where the p values were calculated with edgeR, N=490. (**E**) The subset where the p values were calculated with limma, N=75. (**F**) The subset where the p values were calculated with unassigned analysis platform, N=455. (**G**) Posterior summaries of anti-conservative p value histogram proportions in raw and filtered p value sets. Raw p value data is same as in panel A in S6 Fig., filtered p value data is from simple bernoulli model [anticons ∼ de_tool], N = 3,427. (**H**) Effect size of low-count feature filtering to proportion of anti-conservative p value histograms. (**I**) Posterior summaries of *π*0 values of p value histograms in raw and filtered p value sets. Raw p value data is same as in Fig. 3B and filtered p value data is from beta model [pi0 ∼ de_tool], N = 6,057. (**J**) Effect size of low-count feature filtering to *π*0. The model object related to filtered p value sets in panel G can be downloaded from https://gin.g-node.org/tpall/geo-htseq-paper/raw/26619a4b74aa3781ac6a244edcc24e0ad6eb064b/models/anticons_detool_filtered.rds. The model object related to filtered p value sets in panel I can be downloaded from https://gin.g-node.org/tpall/geo-htseq-paper/raw/26619a4b74aa3781ac6a244edcc24e0ad6eb064b/models/pi0_detool_full_data_filtered.rds.

### Curing of p value histograms by removing low-count features

We observed a small reduction in the mean number of p values per GEO experiment of anti-conservative and uniform histograms as compared to other p value histogram classes, suggesting that p value sets with anti-conservative and uniform shape are more likely to have been pre-filtered, or have been more extensively pre-filtered (S1 Fig.).

Accordingly, we speculated that by further filtering out features with low counts we could convert some of the untoward p value histograms into anti-conservative or uniform types. Our goal was not to provide optimal interventions for individual datasets, which would require tailoring the filtering algorithm for the requirements of a particular experiment, but merely to provide proof of principle evidence for or against the hypothesis that by a simple filtering approach we could increase the proportion of anti-conservative p value sets and/or reduce the dependence of results on the analysis platform. Therefore we applied filtering to 3,550 p value sets where we were able to identify gene expression values (see Methods for details of selecting filtering thresholds). We found that overall we could increase the proportion of anti-conservative p value histograms by 2.4-fold, from 866 (24.4%) to 2,866 (58.8%), and the number of uniform histograms from 9 (0.25%) to 18 (0.5%) (Fig. 4A). For all analysis platforms, most rescued p value distributions came from classes “bimodal” and “other,” while almost no rescue was detected from conservative histograms (Fig 4B-F). Upon removal of low count features the proportion of anti-conservative p value histograms increased for all analysis platforms, with the largest effects observed for cuffdiff and deseq, which presented the lowest pre-rescue fractions of anti-conservative p histograms (Fig 4G, H). Nevertheless, substantial differences between analysis platforms remain, indicating that removal of low-count features, while generally beneficent, was not sufficient to fully remove the sources of bias originating from analysis platform. Also, the *π*_0_-s calculated from the rescued set of anti-conservative p value sets have very similar distributions compared to *π*_0_-s from the pre-rescue anti-conservative p value sets, and concomitantly very similar dependence on the analysis platform (Fig 4I, J).

### Are experiments resulting in anti-conservative p value distributions associated with impact of the resulting publication?

The impact of a scientific paper can be assessed through the impact of the journal that publishes it and on its own merit, as attested by the citations it gathers. We investigated whether the experiments, whose statistical analysis resulted in anti-conservative p value distributions, were published, on average, in higher-performing journals, and whether such papers get more citations. For journal performance we used the Elsevier CiteScore, which employs a relatively long three-year window and all document types published in each journal, making it a potentially more stable and robust metric than journal IF [29,30]. Our time-adjusted analysis shows a negative association (Fig. 5), where in journals with the highest cite scores, in comparisons with the lowest ranked journals, the experiments with anti-conservative p value distributions are underrepresented on average by 10.5 (95% CI 0.34; 17.3) percentage points, and the probability of a positive association is 0.003. This is qualitatively similar to an observed negative association of reproducibility of qPCR experiments with a journal quality metric [31]. A likely reason for this negative association would be the tendency of higher-ranking journals to publish papers with more authors conducting more individual experiments, using a wider variety of experimental systems, potentially leading to less effort and/or competence expended towards individual experiments.

**Fig 5.**
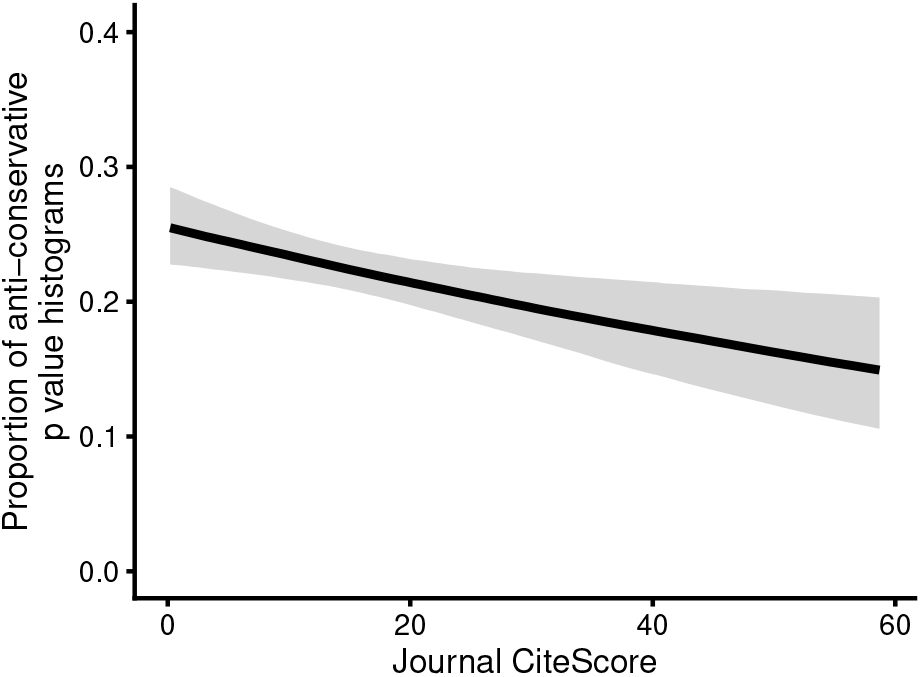
Association of journal CiteScore with proportion of anti-conservative p value histograms. The line denotes best fit of the logistic model [anticons ∼ CiteScore + year, bernoulli likelihood]. Shaded area denotes 95% credible region. N=2,548. The model object can be downloaded from https://gin.g-node.org/tpall/geo-htseq-paper/raw/26619a4b74aa3781ac6a244edcc24e0ad6eb064b/models/anticons__CiteScore_year.rds..

## Discussion

In this work we assess the quality of the differential expression analysis by HT-seq based on a large unbiased NCBI GEO dataset. To the best of our knowledge this is the first large-scale study to offer quantitative insight into general quality of experimentation and data analysis of a large field of biomedical science. We show that

- Overall, three quarters of DE HT-seq experiments result in p value distributions that indicate that assumptions behind the DE tests have not been met.
- There are very few experiments resulting in uniform p value distributions that indicate <100 DE features.
- The distribution of *π*_0_-s, the fraction of true null hypotheses in an experiment, peaks at around 0.5, suggesting that in many experiments most genes are DE.
- The proportion of anti-conservative p value distributions, as well as the values of *π*_0_-s, are associated with the DE analysis platform used.
- For many experiments, a simple exclusion of low-count features rescues the p distribution into anti-conservative form.
- There is a negative association of anti-conservative p value distributions with journal impact factor.

While there is a positive temporal trend for the increasing fraction of anti-conservative p value sets, overall a strong majority of them fall into shapes that indicate that the assumptions behind the statistical tests, which produced these p values, have not been met. Specifically, p value distributions that are not anti-conservative/uniform relinquish statistical control of the experiment over type I errors, on the presence of which later p-value adjustments are predicated [19]. Such p value sets have effectively relinquished their statistical meaning and are thus problematic for scientific interpretation and further analysis, including FDR/q value calculations. Currently most DE analysis platforms use parametric tests accompanied by GLM regression. However, it has been recently shown for the often-used parametric tests that their False Discovery Rate control can be exceedingly poor, while non-parametric Wilcoxon rank sum test achieves better results [32]. Furthermore, it is becoming clear that data normalization can introduce bias into RNA-seq DE analysis, which cannot be corrected by many of the currently widely used methods [33,34]. Indeed, widely used DE analysis tools use RNA-seq data pre-processing strategies, which are vulnerable to the situations where a large fraction of features change expression and/or different samples exhibiting differing total RNA concentrations [23,35]. In general, the analysis tools that are used for differential expression testing allow for a plethora of choices, including distributional models, data transformations and basic analytic strategies, which can lead to different results through different trade-offs [36–38]. Our finding of a high proportion of unruly p value distributions suggests that the wide variety of individual data pre-processing/DE testing workflows, in their actual use by working scientists, results in an overall poor outcome. Although this situation has been steadily improving over the last decade, at least for the limma, edgeR and DESeq users, there remains substantial room for further improvements.

What could cause the systematic differences between analysis platforms that we see in p value distributions and *π*_0_ values? The major points of divergence in the analytic pathway include raw sequence pre-processing, aligning sequences to a reference sequence, counts normalization, and differential expression testing [37,39]. Our inability to abolish analysis platform driven bias by removal of low-count features suggests that pre-filtering of data is not a major source of this bias. Although, in principle any tool can be used in a multitude of workflows, different tools tend to offer their users not only different suggested workflows, but also different amounts of automation to data pre-processing and analysis. Cuffdiff almost fully automates the analysis, including removal of biases [40], DESeq2 workflow suggests some pre-processing steps to speed-up computations but uses automation to remove biases from input data [41], whereas edgeR [42] and limma-voom require more interactive execution of separate pre-processing and analysis steps [43,44]. We speculate that popularity of cuffdiff and DESeq2 partly lies in their automation, as the user is largely relieved from decision making. However, we found that cuffdiff is associated with the smallest proportion of anti-conservative p value histograms, whereas limma and edger, with their more hands-on approach, are associated with the highest proportions of anti-conservative histograms. Interestingly, limma and edgeR use very different distributional models for DE testing, supporting the notion that it might be data pre-processing, rather than the statistical test, that has the most impact on the success of DE analysis [23]. However, limma and edgeR were not the top performers on the *π*_0_-metric, where the highest performance associates with cuffdiff, whose ability to provide anti-conservative p value distributions was by far the least impressive (note that the *π*_0_-s were calculated only from the experiments with well-distributed p values). While most analysed experiments revealed *π*_0_-s far from unity, there were extremely few uniform p value distributions, suggesting relatively few true effects. This unexpected result is made even more surprising by the low statistical power (< 40%) of most real-world RNA-seq DE experiments, which should increase the likelihood of encountering uniform p value distributions [45]. As a technical comment, it should be noted, that the class assigned by us on a p value histogram depends on arbitrarily set bin size. Our use of 40 bins leads to histograms, where an experiment with up to around 100 true effects out of 20 000 features could well lead to a uniform histogram, because of swamping of the lowermost bin (p < 0.025) with p values emanating from true null effects (S18 Fig.). The lack of uniform p value distributions would suggest that (i) there are almost no experiments submitted to GEO, where the experimental treatment in led to only a few DE genes and (ii) that the actual power of GEO-submitted experiments is not low at all. We find both possibilities hard to believe. As for power considerations, we determined from a random sample of 216 GEO submissions that 19% of p value sets were calculated from N = 1 experiments and 50% from N < 3 experiments, whereas only 8% had a sample size of 4 or larger (data not shown). These prevalent sample sizes are clearly incompatible with the literature in terms of providing enough power to the experiment to reliably find most DE genes. For example, with the currently favoured sample size of three, for most genes in isogenic yeast, effect sizes of at least 4-fold seem to be required for successful analysis and the overall minimal acceptable sample size can be from 4 to 6 [45,46]. In animals a reasonable sample size is expected to be substantially larger, over 10 for most genes, which are not highly expressed, and for cancer samples it seems to be well over 20 [38,45,47–49]. This raises the possibility that nearly all experiments, which should have resulted in uniform distributions, were somehow shifted into other distributional classes.

The DE analysis tool-specific means of *π*_0_ values range from 0.5 (tool “unknown”) to 0.7 (tool “cuffdiff”), showing that by this criterion in an average RNA-seq experiment about half of the cellular RNA-s are expected to change their expression levels (Fig. 3B). In principle, a low *π*_0_ could reflect true differential expression as a causal response to experimental treatment, or it could be an artefact of suboptimal data analysis. To further investigate this, we separately analysed two cases of studies related either to cancer or transcription factors, assuming that available p value sets reflect situation where DE is profiled in cancer cells/tissues or after transcription factor interrogation, respectively. Due to large-scale rearrangements in cancer genomes, and ability of many TFs to change expression of many genes, these studies are expected to lead to lower *π*_0_-s, perhaps in the 0.4 range [27]. However, in both cases we saw essentially unchanging *π*_0_ distributions, which suggests that the low *π*_0_-s in our dataset are indicative of problematic analytic workflows. It should be added that as most data normalization workflows assume that most genes are not DE (and that total RNA concentrations are stable across experimental conditions), any study that results in a low *π*_0_ should strive to explicitly address this fact in experimental setup, data analysis, and in interpretation of results. It has been argued that most genome wide DE studies ever conducted, including by HT-seq, have used experimental designs that would make it impossible to untangle such global effects, at least in a quantitatively accurate way [50,51]. The issue lies in the use of internal standards in normalizing the counts for different RNA-s, which leads to wrong interpretations, if most genes change expression in one direction. To overcome this problem, one could use spike-in RNA standards and compositional normalization [35]. However, spike-in normalization requires great care to properly work in such extreme cases [34,52], and outside single-cell RNA-seq it is yet rarely used [35]. Thus, it seems likely that many or most of the low *π*_0_ experiments represent technical failures, most likely during data normalization. When we combine the low power of most RNA-seq DE experiments with our results concerning the near complete lack of uniform p value distributions, the high proportion of improper p value distributions, highly variable and overall quite low *π*_0_-s, and associations with DE analysis platforms, there remains not much room for doubt about existence of pervasive analytic problems in the actual practice of DE RNA-seq. How much this level of technical malpractice influences the quality of the actual scientific conclusions – which is what really matters in the end – is an open question, but one cannot but assume that this influence is substantial, and in need of corrective action. What would this corrective action entail? Currently there exist at least 29 different DE RNA-seq analysis platforms which, among other differences, use 12 different distributional models for DE testing [46]. It is becoming clear that for an average user of RNA-seq a winnowing of recommended analytic choices is needed. Nevertheless, design and analysis of a specific DE RNA-seq experiment would have to be responsive to several factors, including (i) the study organism and its genetic background, pertaining to biological variation and thus sample size; (ii) the expected *π*_0_, pertaining to experimental design, like use of spike-in probes, and to data analysis; (iii) the number of genes of interest, pertaining to sample size through the multiple testing problem; (iv) the expected expression levels of genes of interest and whether DE of gene isoforms is of interest, pertaining to sample size and to sequencing depth; (v) the structure of experiment pertaining to analysis via GLM (is it multi-group, multi-centre, multi-experimenter?). As no analytic workflow has been shown to systematically outperform the others [23,36–38], fool proof best-practices are yet out of reach. Meanwhile, the user is advised to manually incorporate quality control steps into their workflow, like checking p value histograms and *π*_0_-s before advancing to further adjustments. Also, removal of low-expressed genes before model fitting in DE analysis is a recommended step, which can have a substantial positive effect on the sensitivity of detection of differentially expressed genes [53,54]. However, this comes with the cost of excluding about half the genes from analysis as lowly expressed [55]. There is recent evidence for the unreliability of quantification of differential expression of low-expressed genes by existing DE analysis tools [56], and the sample size needed for accurate measurement of DE has a strong inverse relationship with the expression level of the gene [38].

While the aim of this paper was to provide a birds-eye view on the HT-seq field, the dataset created to support this work provides added value by allowing access at individual GEO submission/experiment level. The dataset that accompanies this study allows to get a first estimate for the quality of conclusions of more than 40,000 HT-seq studies deposited in NCBI GEO. The dataset contains for every GEO submission the date of submission, association with publications, organisms studied, association with tabular files; and for every submitted tabular file it contains the name of file, the number of features, the p value histogram type and *π*_0_. Finally, we note that our methodology can be adapted to study any type of experiment that results in at least several hundreds of parallel measurements/p values, such as quantitative protein mass spectroscopy and metabolomics.

## Methods

### NCBI GEO database query and supplementary files

NCBI GEO database queries were performed using Bio.Entrez Python package and by sending requests to NCBI Entrez public API. The exact query string to retrieve GEO HT-seq datasets was ‘expression profiling by high throughput sequencing[DataSet Type] AND (“2000-01-01”[PDAT] : “2020 -12-31”[PDAT]).’ Accession numbers of cancer related datasets were identified by amending original query string with “AND (“neoplasms”[MeSH Terms] OR cancer[All Fields])”. FTP links from GEO datasets document summaries were used to download supplementary file names. Supplementary file names were filtered for downloading, based on file extensions, to keep file names with “tab,” “xlsx,” “diff,” “tsv,” “xls,” “csv,” “txt,” “rtf,” and “tar” file extensions. We dropped the file names where we did not expect to find p values using regular expression “filelist.txt|raw.tar$|readme|csfasta|(big)?wig|bed(graph)?|(broad)?lincs.”

### NCBI supplementary file processing

Downloaded files were imported using Python pandas package, and searched for unadjusted p value sets. Unadjusted p value sets and summarized expression level of associated genomic features were identified using column names. P value columns from imported tables were identified by regular expression “p[^a-zA-Z]{0,4}val,” from these, adjusted p value sets were identified using regular expression “adj|fdr|corr|thresh” and omitted from further analysis. We algorithmically tested the quality of identified p value sets and removed from further analysis apparently truncated or right-skewed sets, p value sets that were not in 0 to 1 range, and p value sets that consisted entirely of NaN values. Columns with expression levels of genomic features were identified by using following regular expressions: “basemean,” “value,” “fpkm,” “logcpm,” “rpkm,” “aveexpr.” Where expression level data were present, raw p values were further filtered to remove low-expression features using following thresholds: basemean=10, logcpm=1, rpkm=1, fpkm=1, aveexpr=3.32. Basemean is a mean of library-size normalized counts of all samples, logcpm is a mean log2 counts per million, rpkm/fpkm is reads/fragments per kilobase of transcript length per million reads, aveexpr is an average expression across all samples, in log2 CPM units, whereas CPM is counts per million. Row means were calculated when there were multiple expression level columns (e.g for each contrast or sample) in table. Filtered p value sets were stored and analysed separately.

### Classification of p value histograms

Raw p value sets were classified based on their histogram shape. Histogram shape was determined based on the presence and location of peaks. P value histogram peaks (bins) were detected using a quality control threshold described in [25], a Bonferroni-corrected alpha-level quantile of the cumulative function of the binomial distribution with size m and probability p. Histograms, where none of the bins were over QC-threshold, were classified as “uniform.” Histograms, where bins over QC-threshold started either from left or right boundary and did not exceeded 1/3 of the 0 to 1 range, were classified as “anti-conservative” or “conservative,” respectively. Histograms with peaks or bumps in the middle or with non-continuous left- or right-side peaks were classified as “other.” Histograms with peaks on both left- and right-side were classified as “bimodal.”

### Calculation of *π*0 statistic

Raw p value sets with anti-conservative shape were used to calculate the *π*0 statistic. The *π*0 statistic was calculated using local FDR method implemented in limma::PropTrueNullByLocalFDR [43] and, independently, Storey’s global FDR smoother method (Storey 2002) as implemented in gdsctools [57] Python package. Differential expression analysis tools were inferred from column names pattern for cuffdiff (column name = “fpkm” and “p value”) [40], DESeq/DESeq2 (column name = “basemean”) [41], EdgeR (column name = “logcpm”) [42], and limma (column name = “aveexpr”) [43], all other unidentified sets were binned as “unknown.”

### Publication data

Publication data were downloaded from NCBI PubMed database using PubMedId-s from GEO document summaries. Journal CiteScore data from years 2011-2019 (October 2020) was downloaded from Elsevier. Yearly journal CiteScore data was merged with GEO series journal publications using journal ISSN/ESSN and articles’ publication year. Sequence read library metadata were downloaded from NCBI SRA database using GEO accessions.

### Modelling

Bayesian modelling was done using R libraries rstan vers. 2.21.3 [58] and brms vers. 1.1.1 [59]. Models were specified using extended R lme4 [60] formula syntax as implemented in R brms package. We used weak priors to fit models. We run minimally 1.1.2 2000 iterations and three chains to fit models. When suggested by brms, Stan NUTS control parameter adapt delta was increased to 0.95–0.99 and max treedepth to 12–15.

### RNA-seq simulation

RNA-seq experiment simulation was done with polyester R package [61] and differential expression was assessed using DESeq2 R package [41] using default settings. Code and workflow used to run and analyze RNA-seq simulations is deposited in Zenodo with doi: 10.5281/zenodo.4463804 (https://doi.org/10.5281/zenodo.4463804). Processed data, raw data and workflow with input fasta file is deposited in Zenodo with doi: 10.5281/zenodo.4463803 (https://doi.org/10.5281/zenodo.4463803).

### Code and raw data

The code to produce raw dataset is available as a snakemake workflow [62] on rstats-tartu/geo-htseq Github repo (https://github.com/rstats-tartu/geo-htseq). Raw dataset produced by the workflow is deposited in Zenodo https://zenodo.org with doi: 10.5281/zenodo.3747112 (https://doi.org/10.5281/zenodo.3747112). The code to produce article’s figures and models is deposited on rstats-tartu/geo-htseq-paper Github repo (https://github.com/rstats-tartu/geo-htseq-paper). Individual model objects are deposited in G-Node with doi: 10.12751/g-node.p34qyd (https://doi.org/10.12751/g-node.p34qyd). ggplot2 vers. 3.3.1[63] R library was used for graphics. Data wrangling was done using tools from tidyverse package [64]. Bayesian models were converted to tidy format and visualised using tidybayes R package [65].

## Acknowledgments

We are grateful for Toomas Mets (University of Tartu) for critically reading the manuscript, and Niilo Kaldalu (University of Tartu) and Margus Pihlak (Tallinn University of Technology) for useful discussions. The work was supported by the European Union from the European Regional Development Fund through the Centre of Excellence in Molecular Cell Engineering (2014-2020.4.01.15-0013) and by the grants from the Estonian Research Council (PRG335, PUT1580).

## Supporting Information

**S1 Fig.**
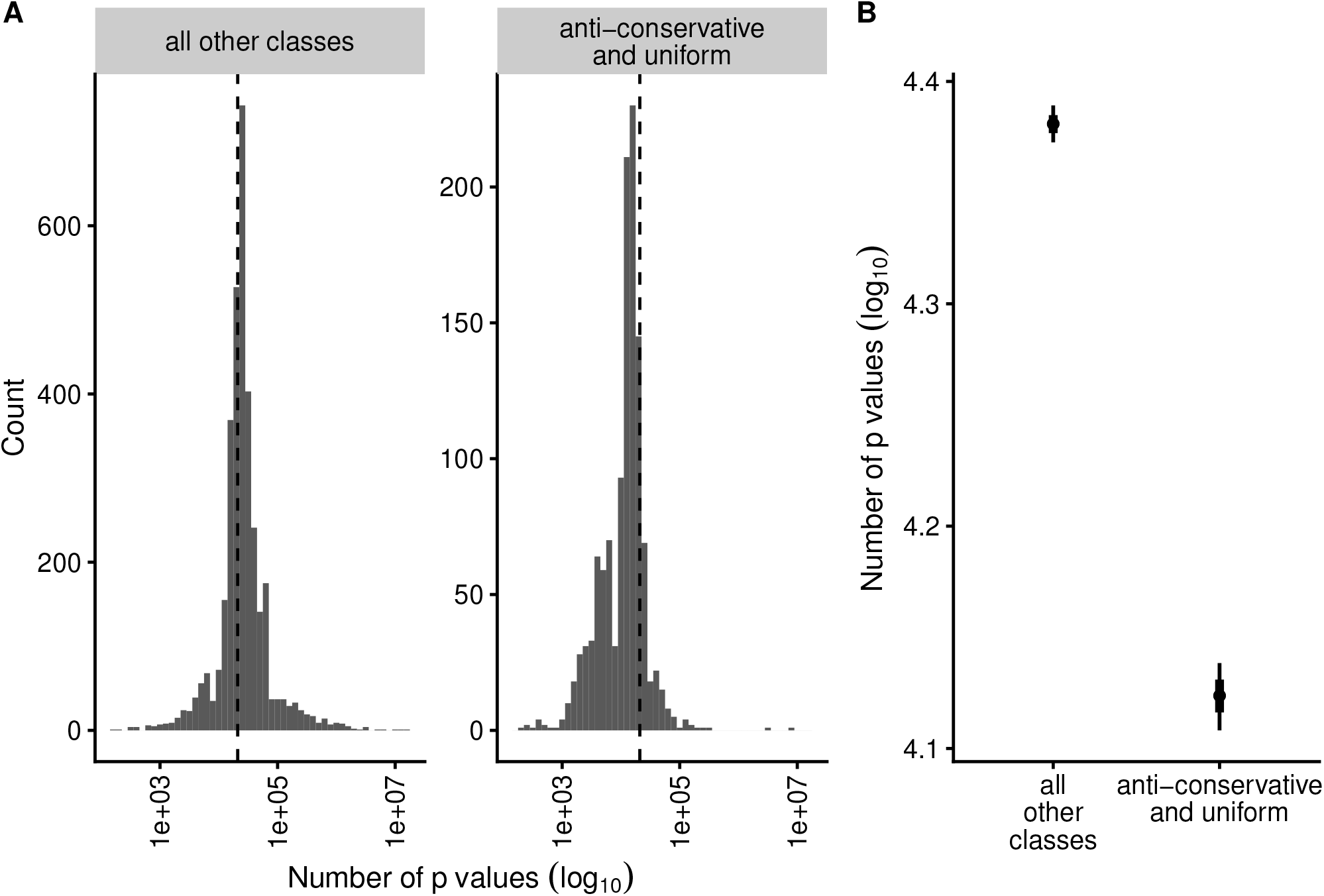
Reduced number of features in anti-conservative and uniform p value sets. (**A**) P value set size distribution. Dashed line denotes the median (20,954) number of features. From each GEO series only one random set was considered, N = 4,616 p value sets. (**B**) Robust linear modeling of number of features in anti-conservative and uniform vs. non-anti-conservative p value sets [log10_n_pvalues ∼ anticons, student’s t likelihood], N = 4,616. Points denote best fit of linear model. Thick and thin lines denote 66% and 95% credible region, respectively. The model object related to panel B can be downloaded from https://gin.g-node.org/tpall/geo-htseq-paper/raw/26619a4b74aa3781ac6a244edcc24e0ad6eb064b/models/log_n_pvalues%20~%20anticons.rds.

**S2 Fig.**
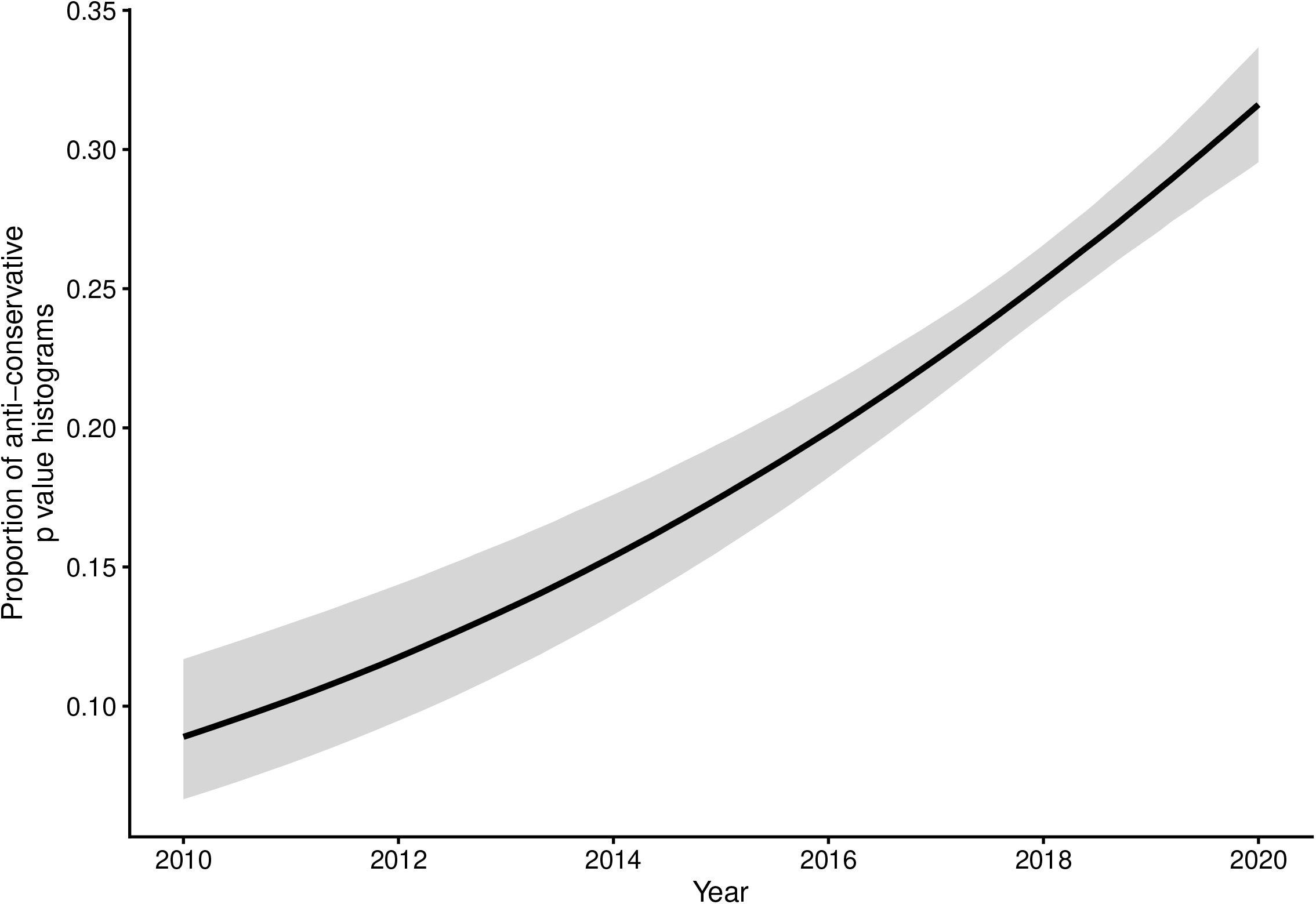
The increasing proportion of anti-conservative histograms. Binomial logistic model [anticons ∼ year], N = 4,616. Lines denote best fit of linear model. Shaded area denotes 95% credible region. The model object related to figure can be downloaded from https://gin.g-node.org/tpall/geo-htseq-paper/raw/26619a4b74aa3781ac6a244edcc24e0ad6eb064b/models/anticons_year.rds.

**S3 Fig.**
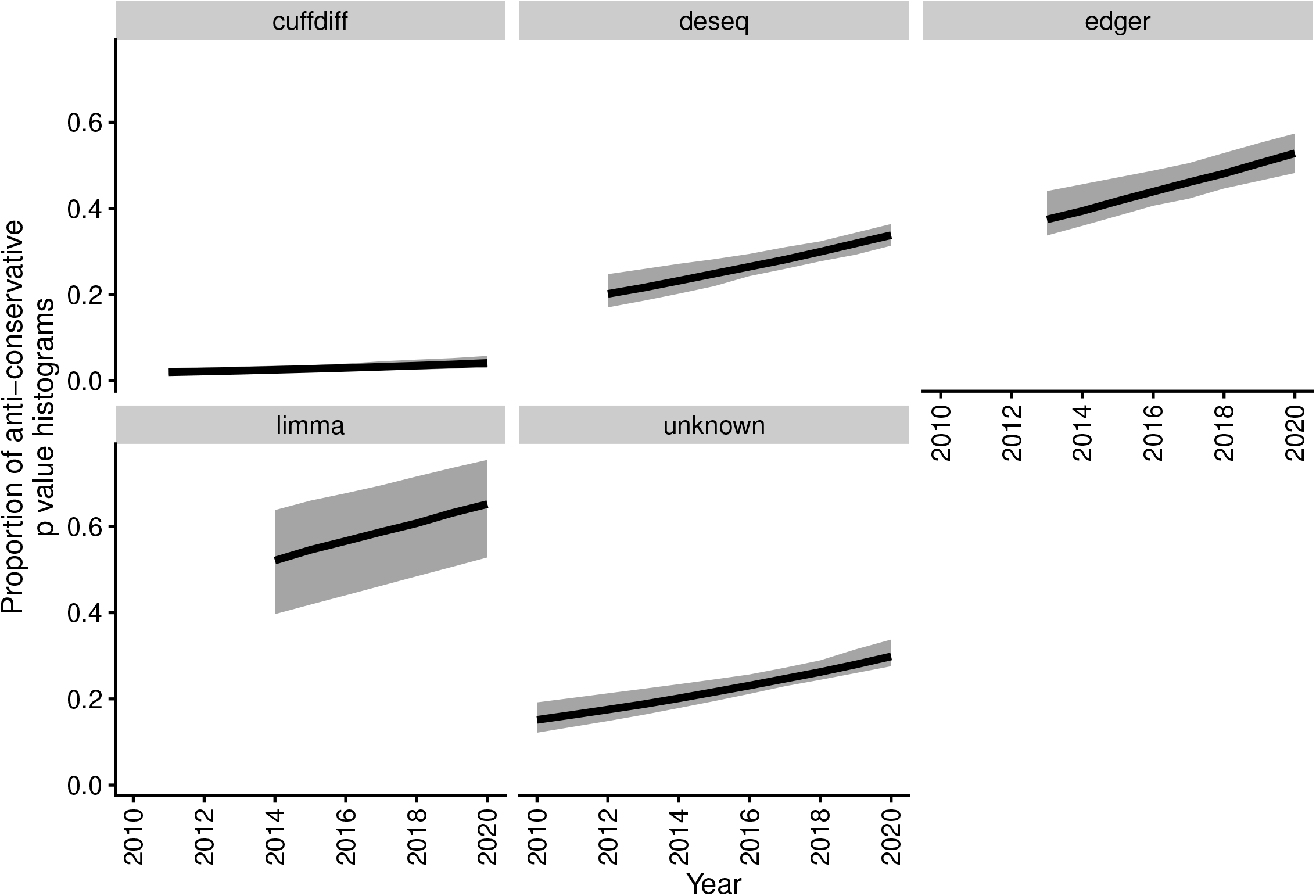
All differential expression analysis tools are associated with temporally increasing anti-conservative p value histograms, two-level binomial logistic model [anticons ∼ year + (year | de_tool)], N = 4,616. Lines denote best fit of linear model. Shaded area denotes 95% credible region. The model object related to figure can be downloaded from https://gin.g-node.org/tpall/geo-htseq-paper/raw/26619a4b74aa3781ac6a244edcc24e0ad6eb064b/models/anticons_year__year_detool.rds.

**S4 Fig.**
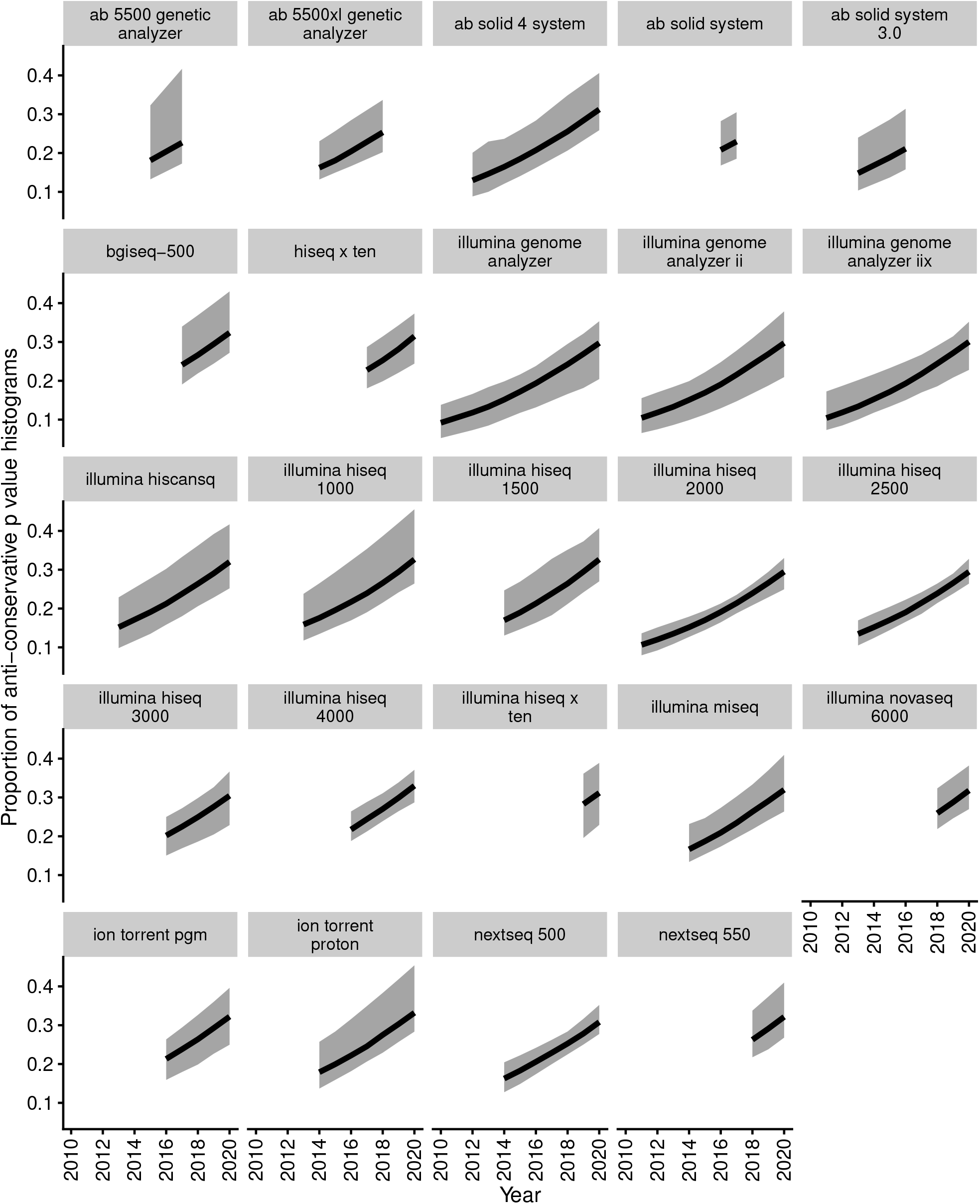
All sequencing instrument models are associated with temporally increasing anti-conservative p value histograms, two-level binomial logistic model [anticons ∼ year + (year | model)], N = 3,778. Only GEO submissions utilizing single sequencing platform were used for model fitting. Lines denote best fit of linear model. Shaded area denotes 95% credible region. The model object related to figure can be downloaded from https://gin.g-node.org/tpall/geo-htseq-paper/raw/26619a4b74aa3781ac6a244edcc24e0ad6eb064b/models/anticons_year__year_model.rds.

**S5 Fig.**
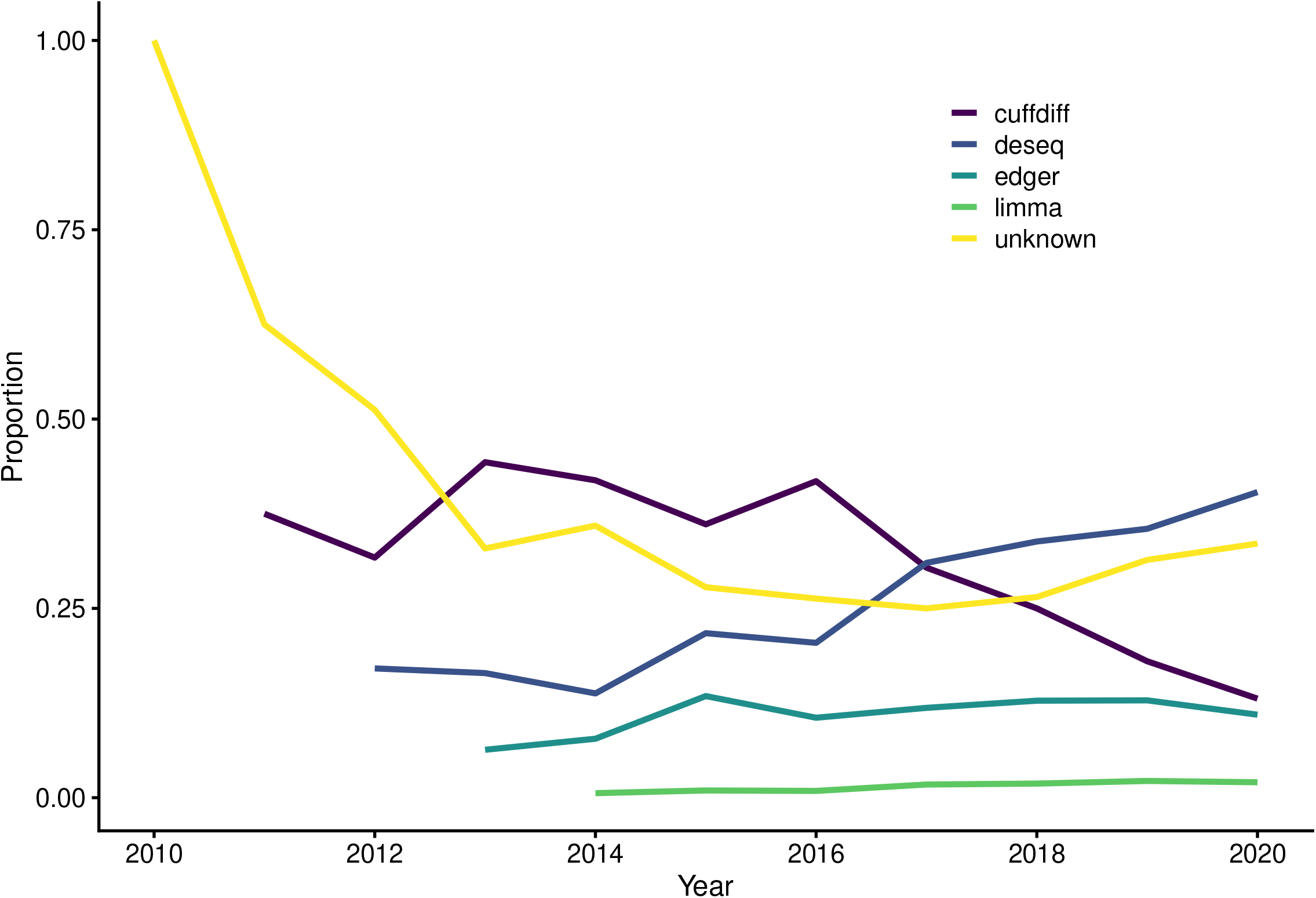
No single differential expression analysis tool dominates the field. Y-axis shows the proportion of analysis platforms, x-axis shows publication year of GEO submission, N = 4,616.

**S6 Fig.**
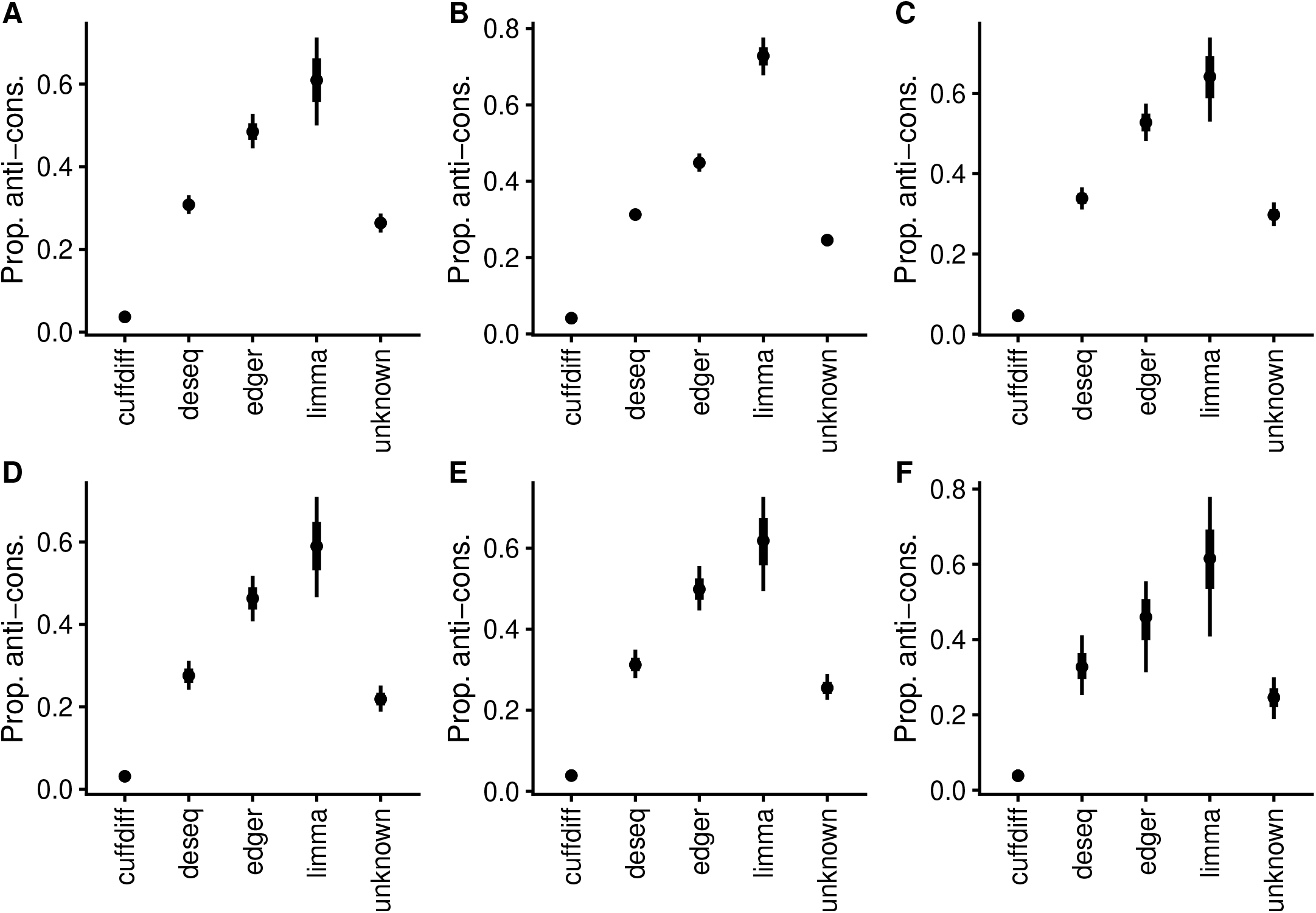
DE analysis tool conditional effects from binomial logistic models for proportion of anti-conservative p value histograms. (**A**) Simple model [anticons ∼ de_tool], N = 4,616. (**B**) Simple model [anticons ∼ de_tool] fitted on complete data, N = 14,813. (**C**) Model conditioned on year of GEO submission [anticons ∼ year + de_tool], N = 4,616. (**D**) Model conditioned on studied organism (human/mouse/other) [anticons ∼ organism + de_tool], N = 3,886. (**E**) Varying intercept model [anticons ∼ de_tool + (1 | model)] where “model” stands for sequencing instrument model, N = 3,778. (**F**) Varying intercept and slope model [anticons ∼ de_tool + (de_tool | model)], N = 3,778. Points denote best fit of linear model. Thick and thin lines denote 66% and 95% credible interval, respectively. The model object related to panel A can be downloaded from https://gin.g-node.org/tpall/geo-htseq-paper/raw/26619a4b74aa3781ac6a244edcc24e0ad6eb064b/models/anticons_detool.rds. The model object related to panel B can be downloaded from https://gin.g-node.org/tpall/geo-htseq-paper/raw/26619a4b74aa3781ac6a244edcc24e0ad6eb064b/models/anticons_detool_all.rds. The model object related to panel C can be downloaded from https://gin.g-node.org/tpall/geo-htseq-paper/raw/26619a4b74aa3781ac6a244edcc24e0ad6eb064b/models/anticons_year_detool.rds. The model object related to panel D can be downloaded from https://gin.g-node.org/tpall/geo-htseq-paper/raw/26619a4b74aa3781ac6a244edcc24e0ad6eb064b/models/anticons_organism_detool.rds. The model object related to panel E can be downloaded from https://gin.g-node.org/tpall/geo-htseq-paper/raw/26619a4b74aa3781ac6a244edcc24e0ad6eb064b/models/anticons_detool__1_model.rds. The model object related to panel F can be downloaded from https://gin.g-node.org/tpall/geo-htseq-paper/raw/26619a4b74aa3781ac6a244edcc24e0ad6eb064b/models/anticons_detool__detool_model.rds.

**S7 Fig.**
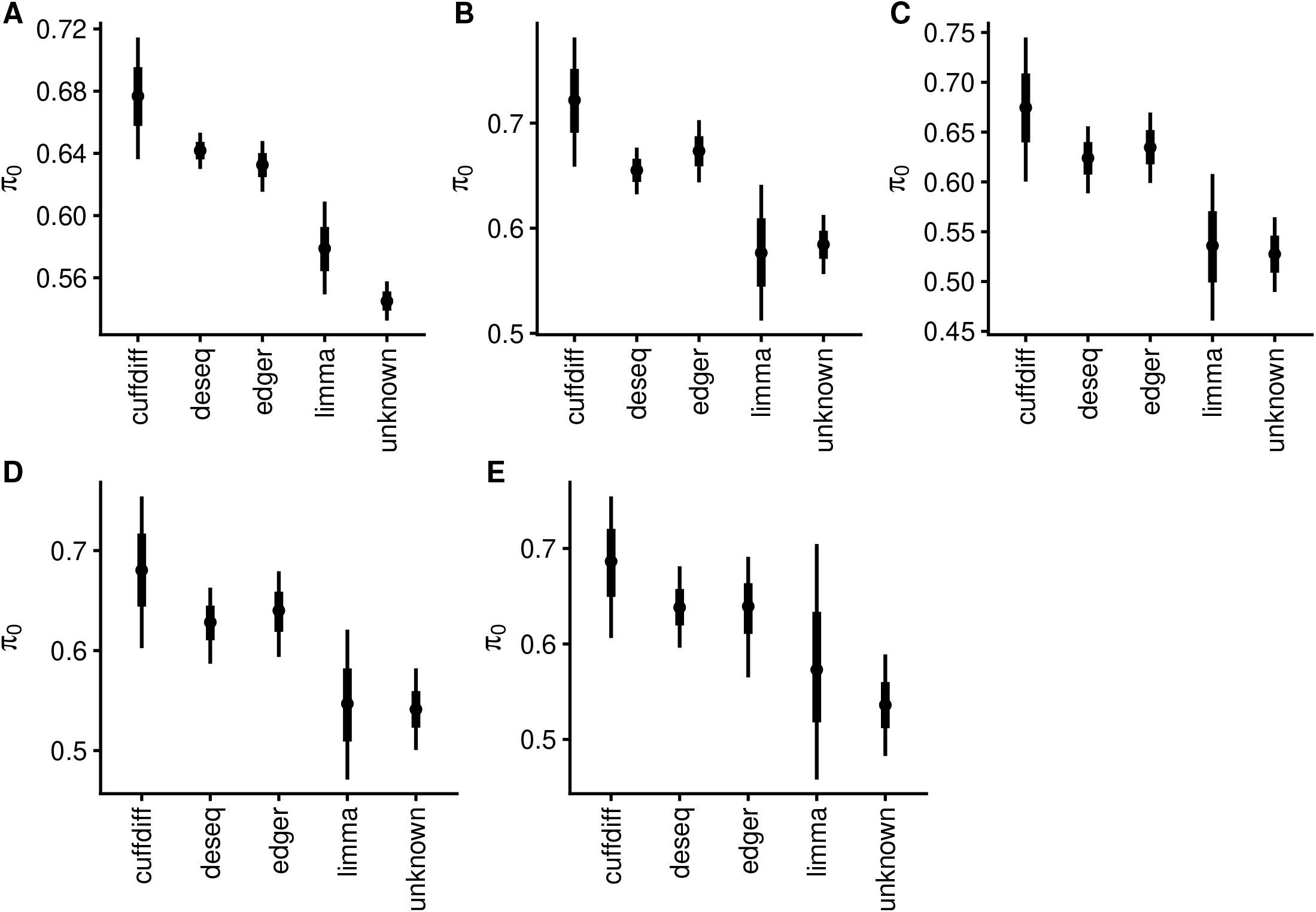
DE analysis tool conditional effects from beta regression modeling of *π*_0_. **A**) Simple model [pi0 ∼ de_tool] fitted on complete data, N = 3,898. (**B**) Model conditioned on year of GEO submission [pi0 ∼ year + de_tool], N = 1,188. (**C**) Model conditioned on studied organism (human/mouse/other) [pi0 ∼ organism + de_tool], N = 993. (**D**) Varying intercept model [pi0 ∼ de_tool + (1 | model)] where ‘model’ stands for sequencing instrument model, N = 959.(**E**) Varying intercept/slope model [pi0 ∼ de_tool + (de_tool | model)], N = 959. Points denote best fit of linear model. Thick and thin lines denote 66% and 95% credible interval, respectively. The model object related to panel A can be downloaded from https://gin.g-node.org/tpall/geo-htseq-paper/raw/26619a4b74aa3781ac6a244edcc24e0ad6eb064b/models/pi0_detool_full_data.rds. The model object related to panel B can be downloaded from https://gin.g-node.org/tpall/geo-htseq-paper/raw/26619a4b74aa3781ac6a244edcc24e0ad6eb064b/models/pi0_year_detool.rds. The model object related to panel C can be downloaded from https://gin.g-node.org/tpall/geo-htseq-paper/raw/26619a4b74aa3781ac6a244edcc24e0ad6eb064b/models/pi0_organism_detool.rds. The model object related to panel D can be downloaded from https://gin.g-node.org/tpall/geo-htseq-paper/raw/26619a4b74aa3781ac6a244edcc24e0ad6eb064b/models/pi0_detool__1_model.rds. The model object related to panel E can be downloaded from https://gin.g-node.org/tpall/geo-htseq-paper/raw/26619a4b74aa3781ac6a244edcc24e0ad6eb064b/models/pi0_detool__detool_model.rds.

**S8 Fig.**
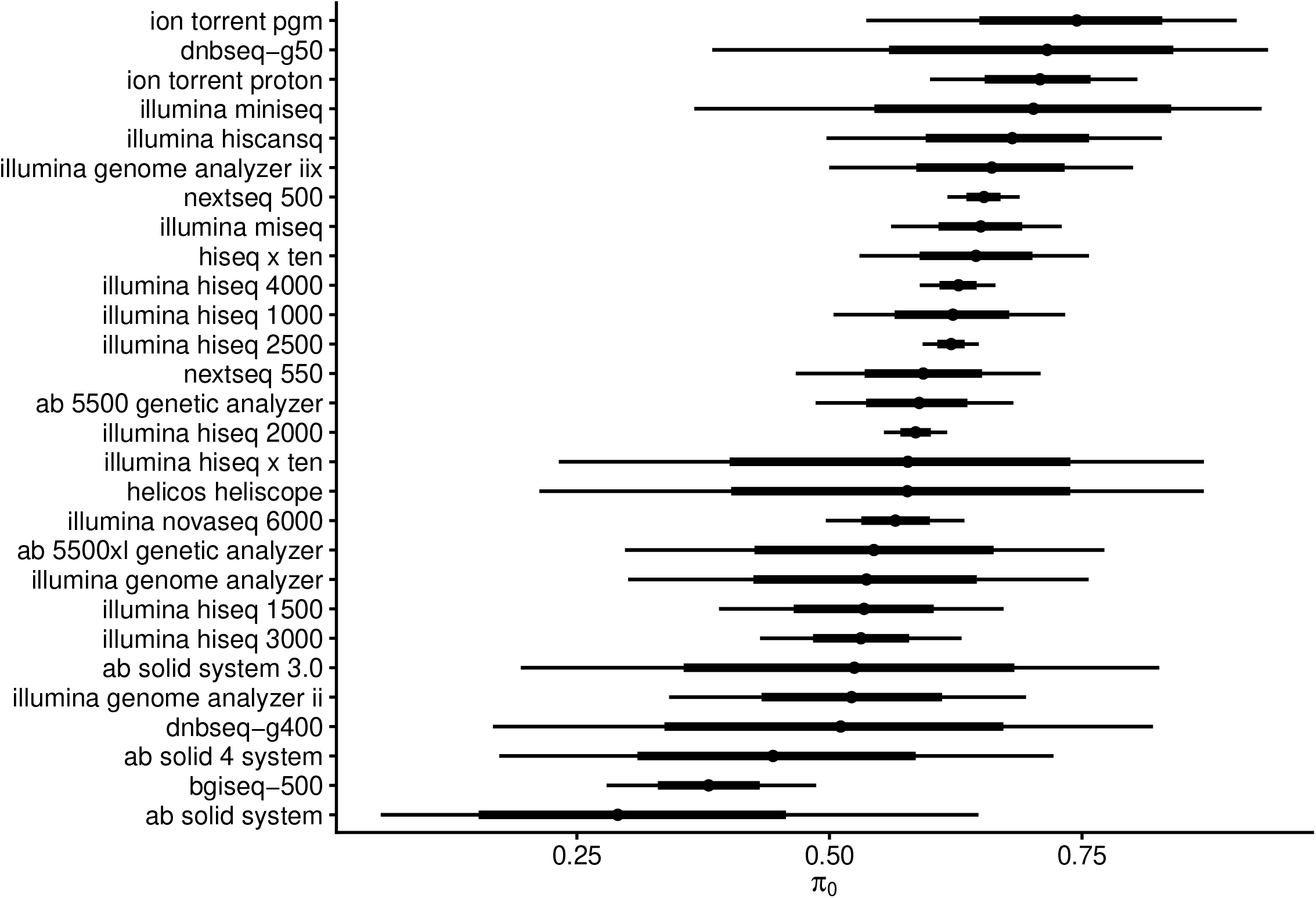
Modeling dependency of *π*_0_ on sequencing instrument model. [pi0 ∼ model], beta distribution, N = 959. Points denote best fit of linear model. Thick and thin lines denote 66% and 95% credible interval, respectively. The model object related to figure can be downloaded from https://gin.g-node.org/tpall/geo-htseq-paper/raw/26619a4b74aa3781ac6a244edcc24e0ad6eb064b/models/pi0__model.rds.

**S9 Fig.**
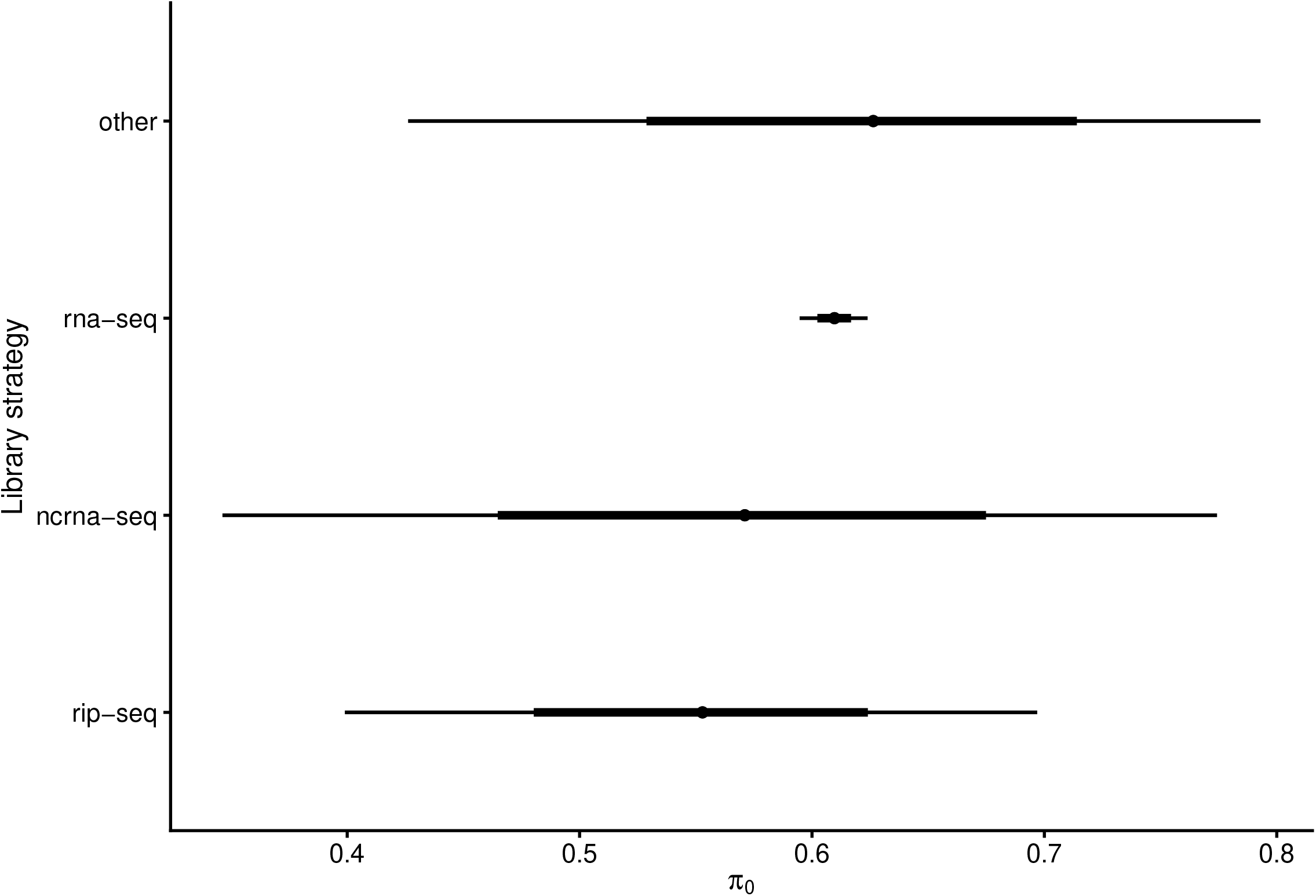
Modeling dependency of *π*_0_ on library strategy. [pi0 ∼ library_strategy], beta distribution, N = 959. Points denote best fit of linear model. Thick and thin lines denote 66% and 95% credible interval, respectively. The model object related to figure can be downloaded from https://gin.g-node.org/tpall/geo-htseq-paper/raw/26619a4b74aa3781ac6a244edcc24e0ad6eb064b/models/pi0__librarystrategy.rds.

**S10 Fig.**
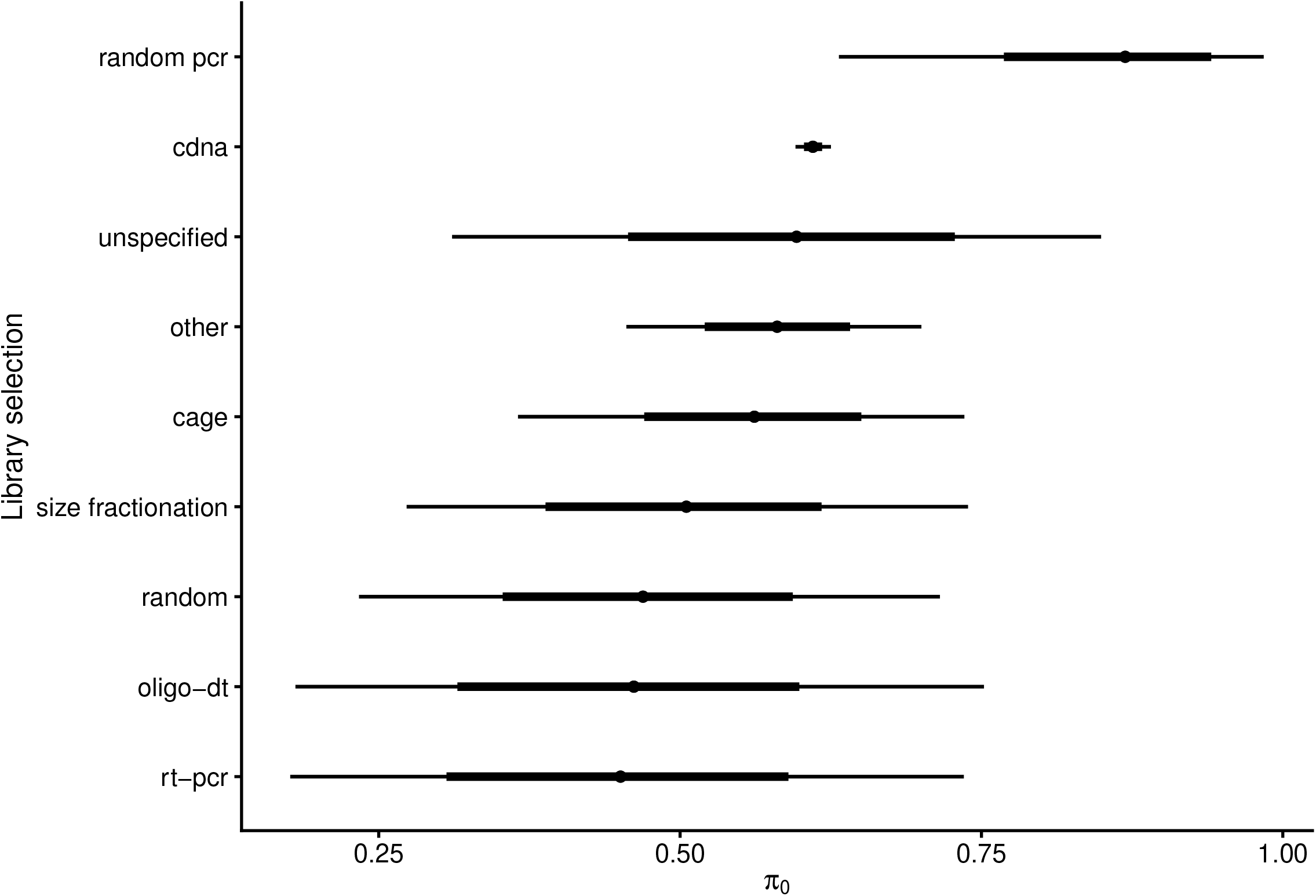
Modeling dependency of *π*_0_ on library selection. [pi0 ∼ library_selection, beta likelihood], N = 959. Points denote best fit of linear model. Thick and thin lines denote 66% and 95% credible interval, respectively. The model object related to figure can be downloaded from https://gin.g-node.org/tpall/geo-htseq-paper/raw/26619a4b74aa3781ac6a244edcc24e0ad6eb064b/models/pi0__libraryselection.rds.

**S11 Fig.**
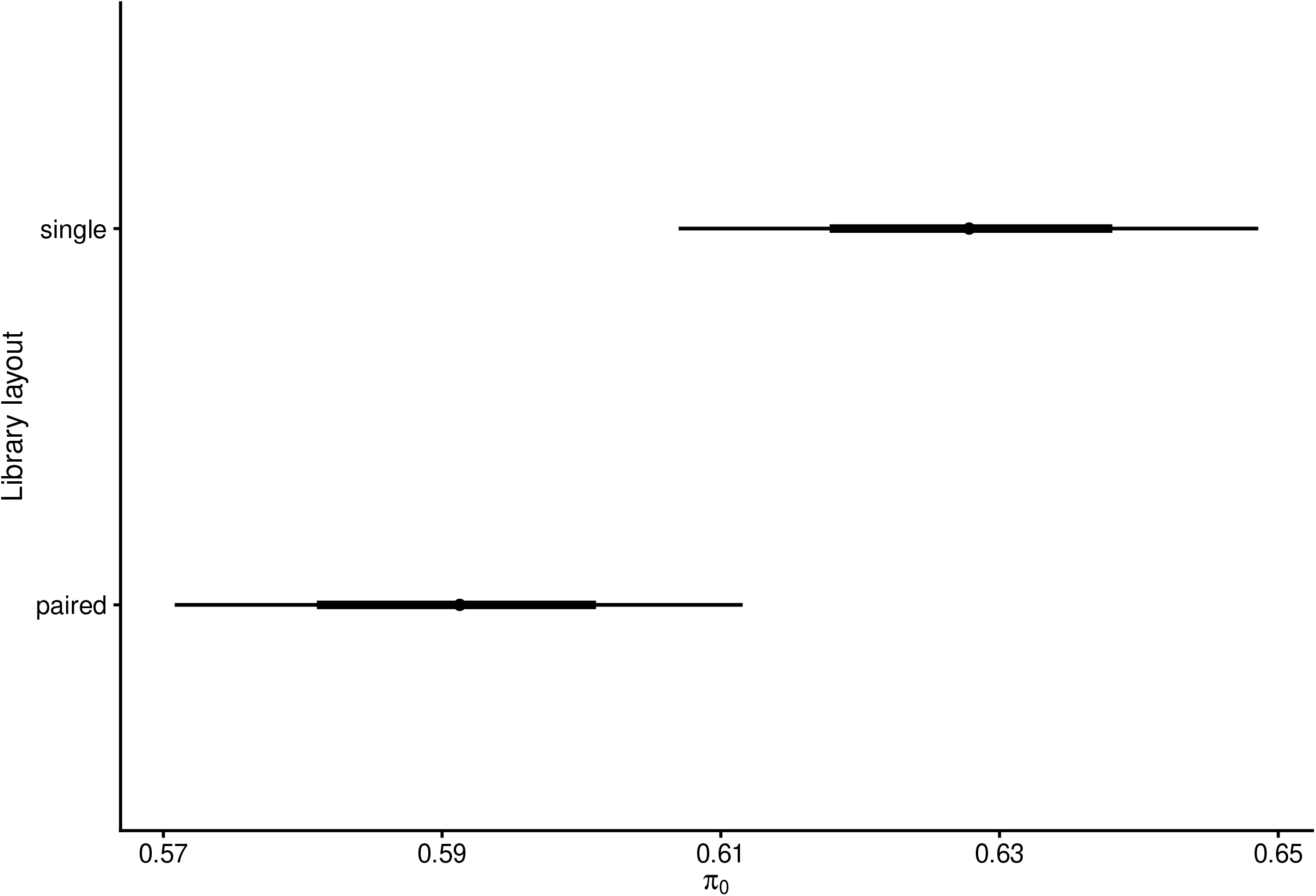
Modeling dependency of *π*_0_ on library layout. [pi0 ∼ library_layout, beta likelihood], N = 959. Points denote best fit of linear model. Thick and thin lines denote 66% and 95% credible interval, respectively. The model object related to figure can be downloaded from https://gin.g-node.org/tpall/geo-htseq-paper/raw/26619a4b74aa3781ac6a244edcc24e0ad6eb064b/models/pi0__librarylayout.rds.

**S12 Fig.**
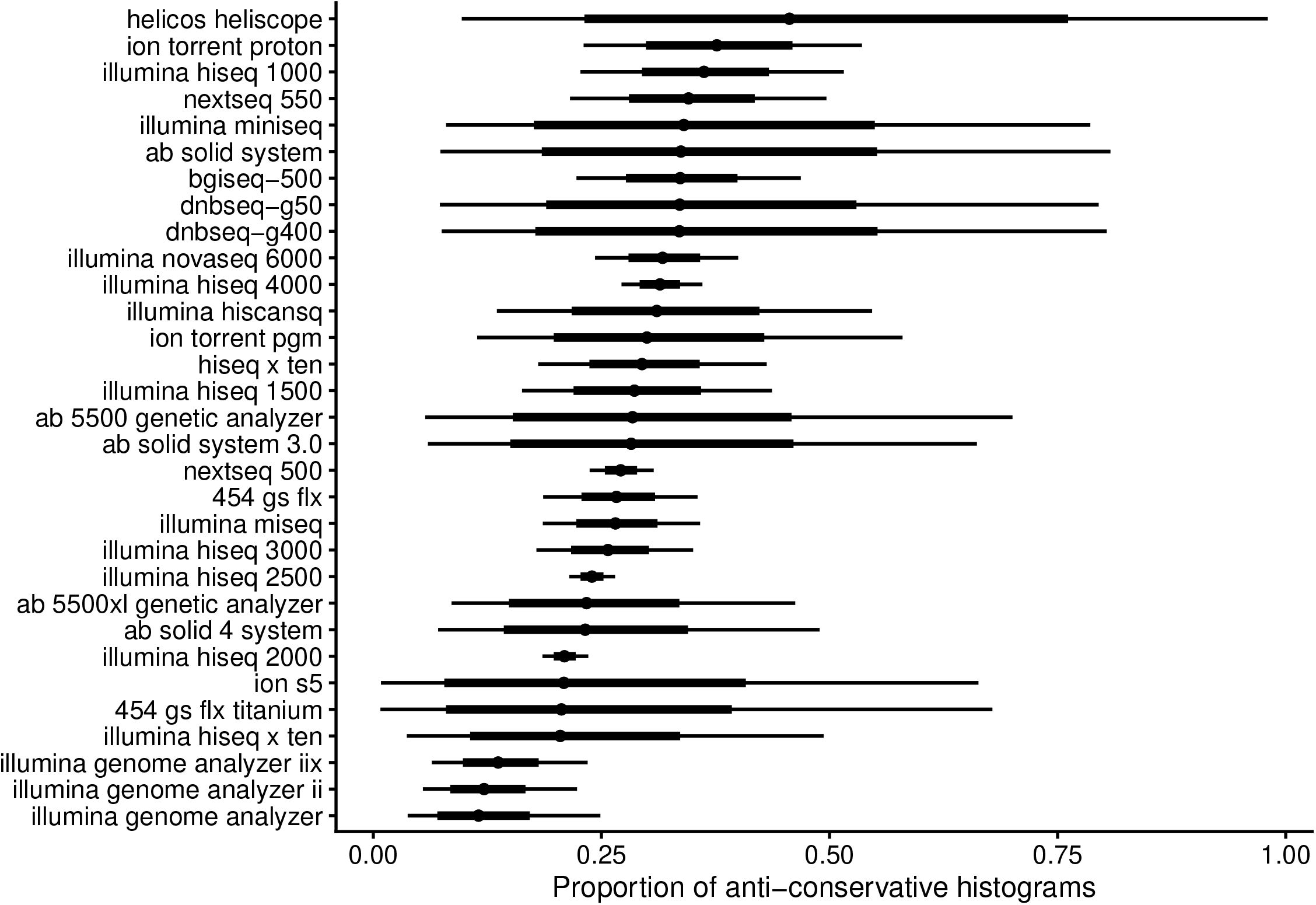
Modeling dependency of proportion of anti-conservative histograms on sequencing platform. [anticons ∼ model, bernoulli likelihood], N = 3,778. Points denote best fit of linear model. Thick and thin lines denote 66% and 95% credible interval, respectively. The model object related to figure can be downloaded from https://gin.g-node.org/tpall/geo-htseq-paper/raw/26619a4b74aa3781ac6a244edcc24e0ad6eb064b/models/anticons__model.rds.

**S13 Fig.**
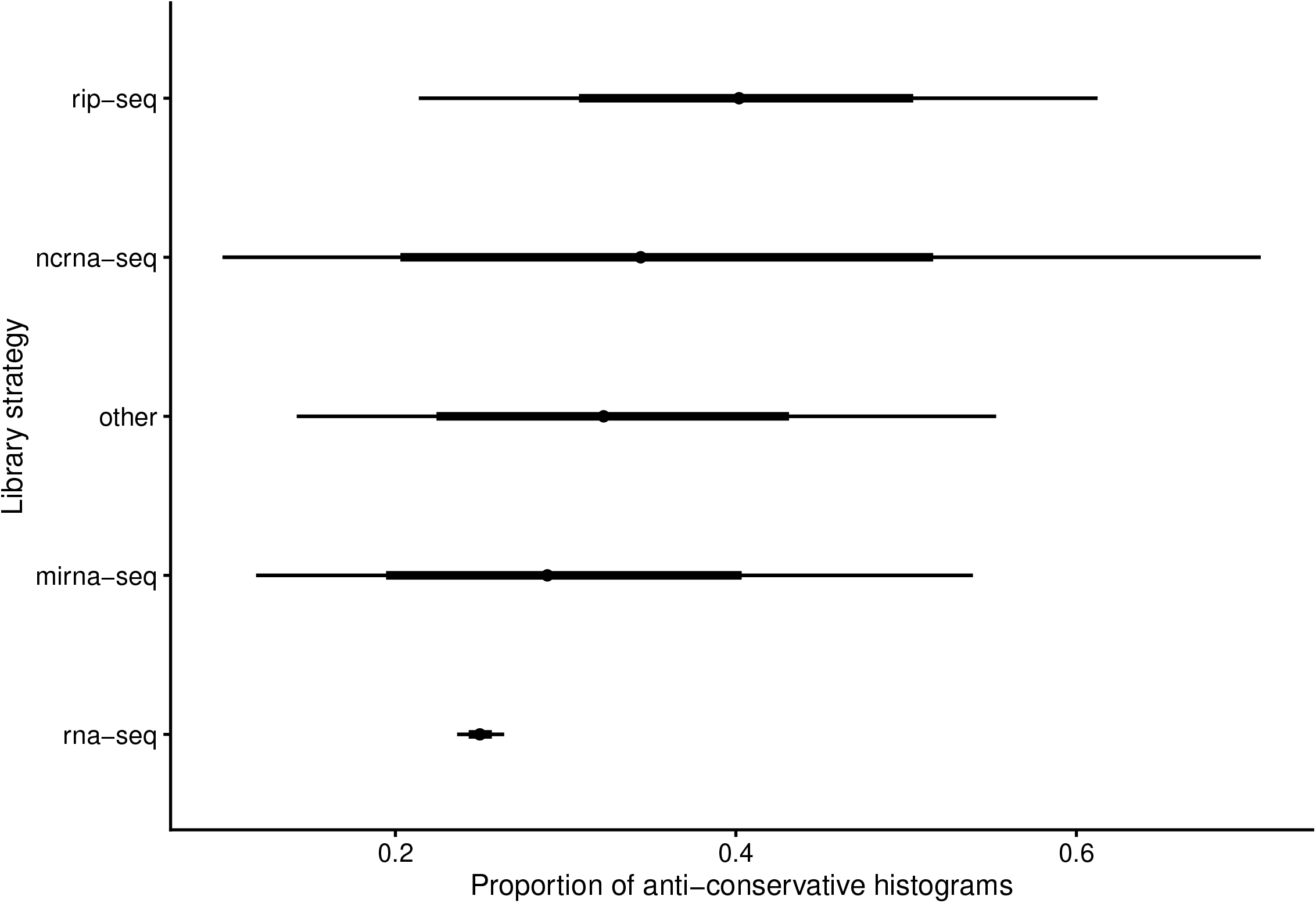
Modeling dependency of proportion of anti-conservative histograms on library strategy. [anticons ∼ library_strategy, bernoulli likelihood], N = 3,778. Points denote best fit of linear model. Thick and thin lines denote 66% and 95% credible interval, respectively. The model object related to figure can be downloaded from https://gin.g-node.org/tpall/geo-htseq-paper/raw/26619a4b74aa3781ac6a244edcc24e0ad6eb064b/models/anticons__librarystrategy.rds.

**S14 Fig.**
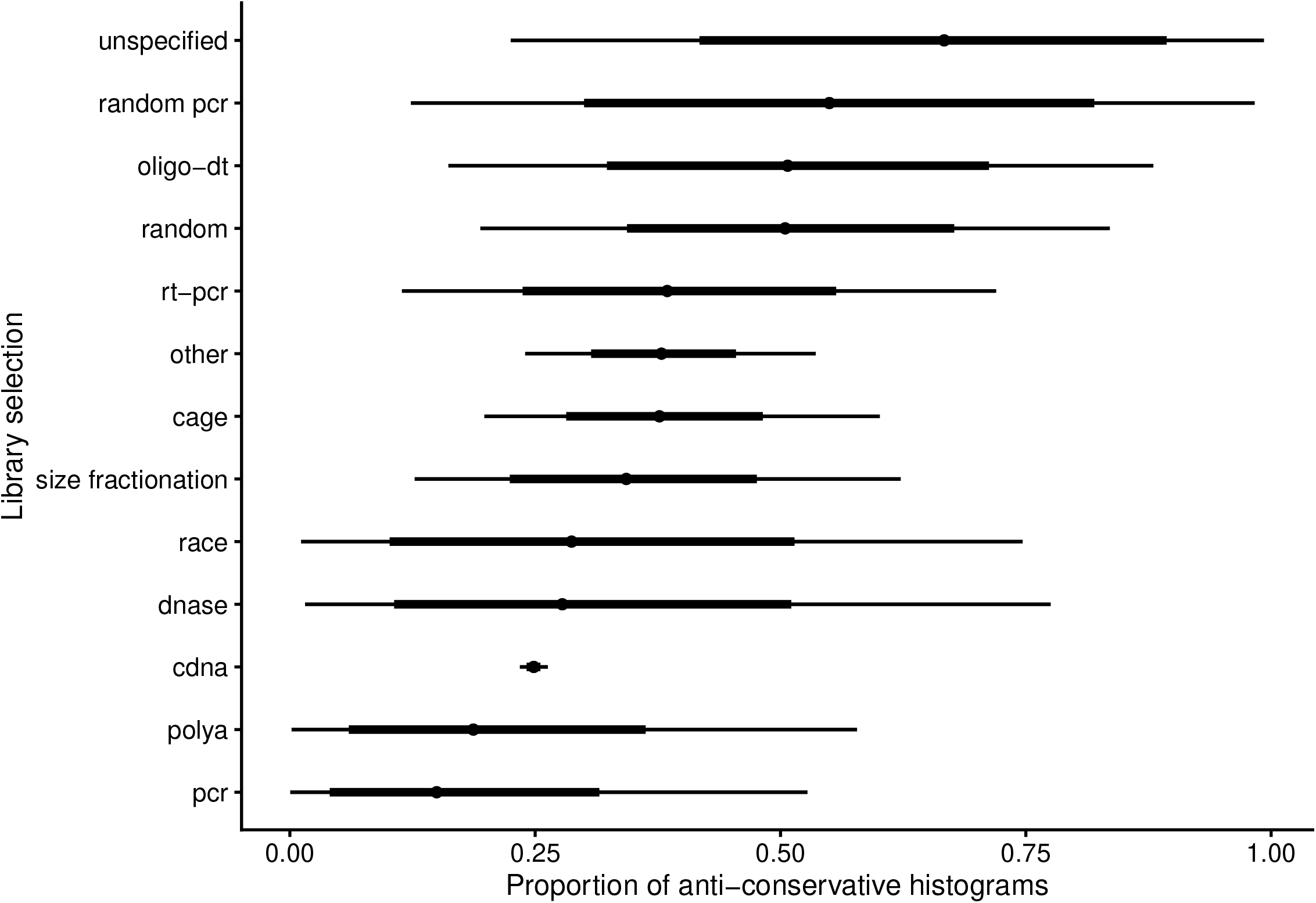
Modeling dependency of proportion of anti-conservative histograms on library selection. [anticons ∼ library_selection, bernoulli likelihood], N = 3,778. Points denote best fit of linear model. Thick and thin lines denote 66% and 95% credible interval, respectively. The model object related to figure can be downloaded from https://gin.g-node.org/tpall/geo-htseq-paper/raw/26619a4b74aa3781ac6a244edcc24e0ad6eb064b/models/anticons__libraryselection.rds.

**S15 Fig.**
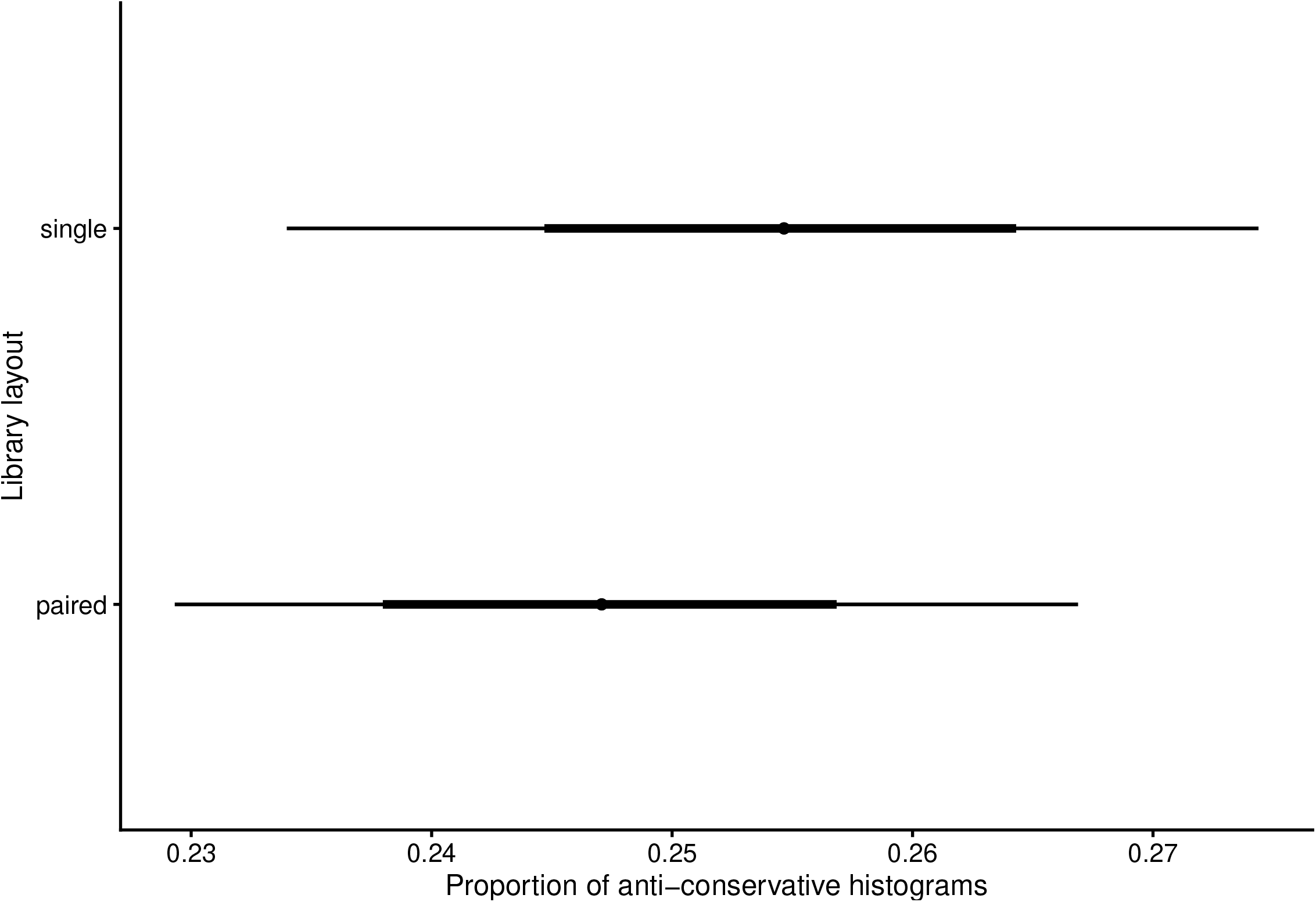
Modeling dependency of proportion of anti-conservative histograms on library layout. [anticons ∼ library_layout, bernoulli likelihood], N = 3,778. Points denote best fit of linear model. Thick and thin lines denote 66% and 95% credible interval, respectively. The model object related to figure can be downloaded from https://gin.g-node.org/tpall/geo-htseq-paper/raw/26619a4b74aa3781ac6a244edcc24e0ad6eb064b/models/anticons__librarylayout.rds.

**S16 Fig.**
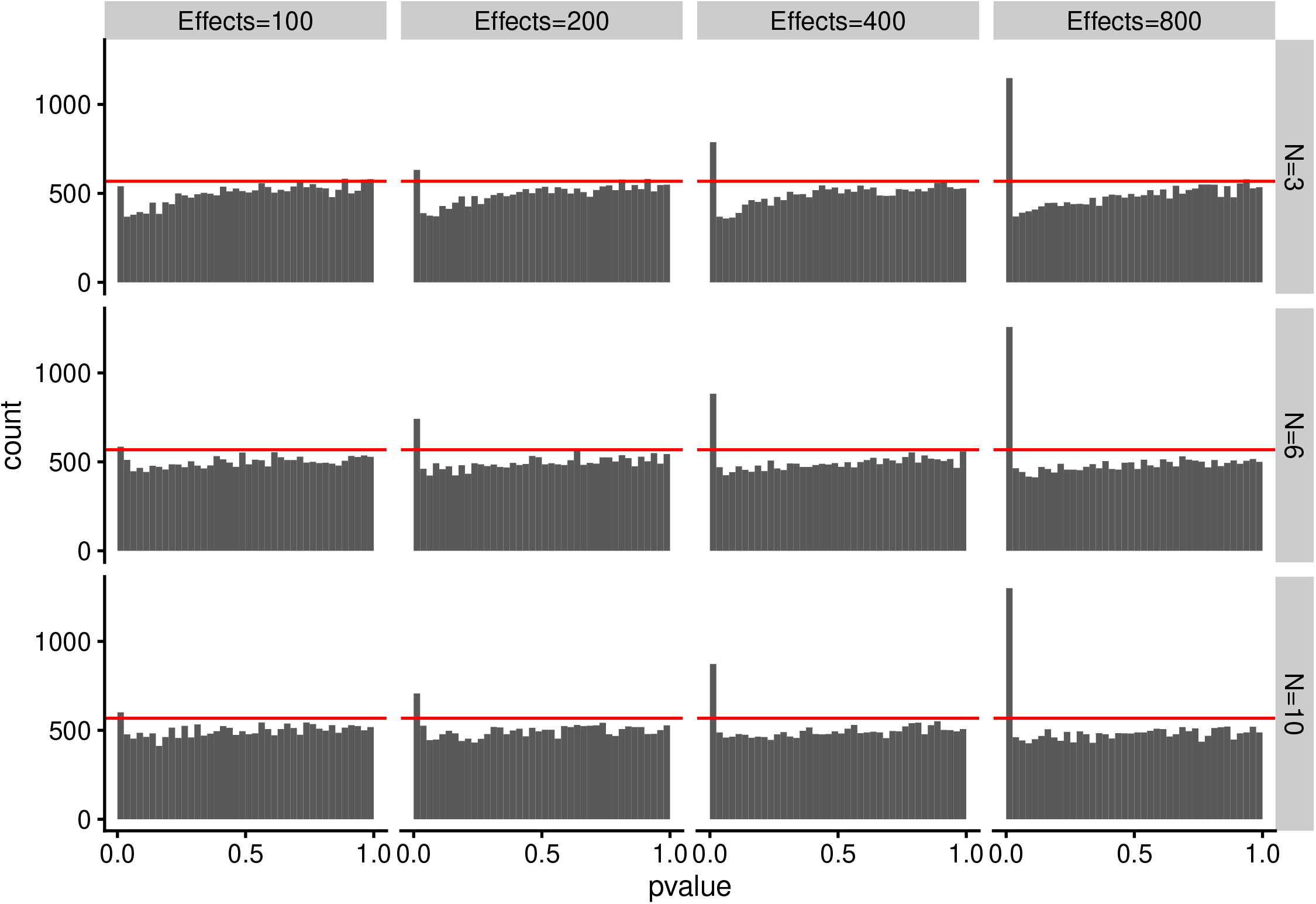
Simulated RNA-seq data shows that histograms from p value sets with around one hundred true effects out of 20,000 features can be classified as “uniform”. RNA-seq data was simulated with polyester R package (Frazee et al. 2015) on 20,000 transcripts from human transcriptome using grid of 3, 6, and 10 replicates and 100, 200, 400, and 800 effects for two groups. Fold changes were set to 0.5 and 2. Differential expression was assessed using DESeq2 R package (Love, Huber, and Anders 2014) using default settings and group 1 versus group 2 contrast. Effects denotes in facet labels the number of true effects and N denotes number of replicates. Red line denotes QC threshold used for dividing p histograms into discrete classes. Code and workflow used to run these simulations is available on Github: https://github.com/rstats-tartu/simulate-rnaseq. Raw data of the figure is available on Zenodo https://zenodo.org with doi: 10.5281/zenodo.4463803.

